# CTCF depletion decouples enhancer-mediated gene activation from chromatin hub formation during cellular differentiation

**DOI:** 10.1101/2024.11.20.624447

**Authors:** Magdalena A. Karpinska, Yi Zhu, Zahra Fakhraei Ghazvini, Shyam Ramasamy, Mariano Barbieri, T. B. Ngoc Cao, Natalie Varahram, Abrar Aljahani, Michael Lidschreiber, Argyris Papantonis, A. Marieke Oudelaar

## Abstract

Enhancers and promoters interact in 3D chromatin structures to regulate gene expression. Here, we characterize the mechanisms that drive the formation of these structures and their function in gene regulation in a lymphoid-to-myeloid transdifferentiation system. Based on analyses at base-pair resolution, we demonstrate a close correlation between binding of regulatory proteins, formation of chromatin interactions, and gene expression. Integration of multi-way interaction analyses and computational modeling shows that tissue-specific gene loci are organized into chromatin hubs, characterized by cooperative interactions between multiple enhancers, promoters, and CTCF-binding sites. Depletion of CTCF strongly impairs the formation of these structures. However, the effects of CTCF depletion on gene expression are modest and can be explained by rewired enhancer-promoter interactions. This demonstrates an instructive role for enhancer-promoter interactions in gene regulation that is independent of cooperative interactions in chromatin hubs. Together, these results contribute to a mechanistic understanding of the structure-function relationship of the genome during cellular differentiation.

## INTRODUCTION

Precise spatio-temporal regulation of gene expression during differentiation and development is dependent on the *cis*-regulatory elements of the genome, which include enhancers and promoters. Active enhancers recruit transcription factors and coactivators, which stimulate assembly and activation of the transcription machinery at gene promoters^1,2^. Since mammalian enhancers can be separated by large genomic distances from their target gene promoters, enhancers interact with promoters in three-dimensional (3D) chromatin structures to activate gene expression^3–7^. Interactions between enhancers and promoters predominantly occur within Topologically Associating Domains (TADs)^8^. TADs are relatively insulated regions of the genome that are demarcated by CTCF-binding sites (CBSs) and formed by loop extrusion^9,10^. During this process, cohesin complexes reel in the chromatin fiber and thereby form progressively larger loops, until they encounter CTCF molecules^11,12^. Although the general importance of loop extrusion for gene regulation remains unclear, it has been shown that loop extrusion contributes to the formation and specificity of (long-range) enhancer-promoter interactions in some contexts^13,14^. In addition, affinity between transcription factors and coactivators bound at enhancers and promoters is thought to have a role in the formation of specific interactions between these elements^15,16^. It has been suggested that these interactions form in the context of (phase-separated) nuclear condensates, which are dependent on multivalent interactions between intrinsically disordered regions of transcription factors, coactivators, the transcription machinery, and/or RNA molecules^17–20^; this model remains debated, however, as it is difficult to test *in vivo*^21^.

Although our understanding of the 3D structures into which the genome is organized has advanced over the last decades, the relationship between genome structure and function remains unclear. In particular, it is incompletely understood when and how enhancer-promoter interactions form during cellular differentiation and how they influence gene expression. The structure-function relationship of the genome has been studied in several developmental contexts, including early embryonic development^22–25^, lineage specification^26,27^, limb development^28^, erythroid differentiation^29^, neuronal differentiation^30,31^, cardiac differentiation^32^, macrophage differentiation^33^, adipocyte differentiation^34^, epidermal differentiation^35^, and cellular reprogramming^36,37^. These studies have identified both instructive enhancer-promoters interactions, which co-occur with active gene expression, and permissive interactions, which are formed prior to gene activation^38^. However, since these studies are based on relatively low-resolution analyses of interactions between enhancers and promoters and do not take their 3D configuration in higher-order structures into consideration, the precise relationship between enhancer-promoter interactions and gene activation, as well as the underlying molecular mechanisms, remain poorly understood. In addition, it is not clear whether permissive, pre-formed interactions represent a distinct mode of enhancer-mediated gene activation or reflect instructive enhancer-promoter interactions that are dependent on subtle changes during cellular differentiation that cannot be detected with low-resolution analyses. Importantly, it has recently been demonstrated that small changes in genome structure can have a big effect of gene expression^39,40^. A better understanding of the structure-function relationship of the genome during cellular differentiation therefore requires analysis at very high resolution and sensitivity. In addition, the integration of perturbations during differentiation could facilitate causal inference and identification of the molecular mechanisms involved.

The analysis of enhancer-promoter interactions in the above-mentioned studies is based on Chromosome Conformation Capture (3C) techniques, which rely on digestion and subsequent proximity ligation of crosslinked chromatin to detect spatial proximity between DNA sequences^41^. The resolution of 3C predominantly depends on the digestion and sequencing strategy^42^. By combining digestion with micrococcal nuclease (MNase), which cuts the genome largely independent of DNA sequence, with deep, targeted sequencing of multiplexed viewpoints of interest, the recently developed Micro-Capture-C (MCC) method supports 3C analysis at single base-pair resolution^43^. Despite their superior resolution, MCC data cannot resolve how *cis*-regulatory elements interact together in higher-order 3D chromatin structures, because MCC is based on the detection of pair-wise interactions in a cell population and therefore cannot distinguish simultaneous, cooperative interactions in individual cells from mutually exclusive interactions that occur independently in different cells. 3D relationships between multiple *cis-*regulatory elements can be disentangled by multi-way 3C techniques, in which multiple interactions derived from individual alleles are captured within (relatively) long sequencing reads^44–49^, and by ligation-free techniques based on physical separation and labeling of crosslinked chromatin interactions^50,51^. Among these, the Tri-C method provides an effective approach to study multi-way enhancer-promoter interactions during cellular differentiation, since it allows for targeted analysis of multiplexed viewpoints of interest at a relatively high resolution (500-5000 bp)^44^. So far, Tri-C and other targeted multi-way 3C techniques have only been used to study a few genetic loci and have not been combined with perturbations of regulatory proteins to study the mechanisms underlying multi-way chromatin interactions. In addition, neither MCC nor Tri-C have been applied throughout subsequent stages of differentiation and development.

In this study, we characterize the structure-function relationship of the genome during lymphoid-to-myeloid transdifferentiation in detail. To this end, we combine analysis of nascent gene expression, chromatin accessibility, binding of regulatory proteins, and chromatin interactions detected by MCC and Tri-C. Integrative analyses of these data show the dynamic formation and dissolution of pair-wise and cooperative multi-way interactions between enhancers, promoters and CBSs during cellular differentiation. Further experiments after auxin-induced CTCF depletion and computational modeling show that CTCF is not required for the formation of pair-wise enhancer-promoter interactions, but essential for the formation of chromatin hubs during cellular differentiation. CTCF depletion therefore allows us to separate the function of pair-wise enhancer-promoter interactions and chromatin hubs. With this approach, we demonstrate that gene regulation during cellular differentiation is instructed by pair-wise enhancer-promoter interactions, but not dependent on cooperative interactions between multiple enhancers and promoters in CTCF-dependent chromatin hubs. Together, these results provide important new insights into the relationship between genome structure and function during cellular differentiation.

## RESULTS

### Chromatin architecture through lymphoid-to-myeloid transdifferentiation

To study the structure-function relationship of the genome, we characterized several structural and functional genomic features during a lymphoid-to-myeloid transdifferentiation trajectory. To this end, we used a B-cell leukemia cell line (BLaER1) that can be efficiently converted into functional induced macrophages (iMacs) by exogenous expression of the transcription factor CCAAT enhancer-binding protein alpha (CEBPA) over the course of 168 h^52^ (**Fig. 1a**). Characterization of this system by qPCR and FACS shows that cellular transdifferentiation occurs synchronously and completely, with approximately 90% of B-cells converting into iMacs over the differentiation course (**Extended Data Fig. 1a,b**). Previous studies have analyzed nascent RNA synthesis by TT-seq^53^; chromatin accessibility by ATAC-seq^53^; the distribution of H3K27ac, H3K27me3, and CTCF by ChIP-seq^54^; and genome organization by Hi-C^54^ during BLaER1 transdifferentiation. We complemented these data by mapping the binding profiles of Mediator and cohesin using an optimized ChIPmentation protocol^55^ at 0 h, 12 h, 24 h, 72 h, and 96 h after differentiation induction (Methods; later timepoints were omitted, as iMacs are quiescent). In addition, we generated Capture-C and MCC interaction profiles from the viewpoints of 51 promoters of B-cell-specific and iMac-specific genes at 0 h, 24 h, and 96 h (**Fig. 1b-e, Extended Data Fig. 1c-e, and Supplementary Table 1**).

**Fig. 1:**
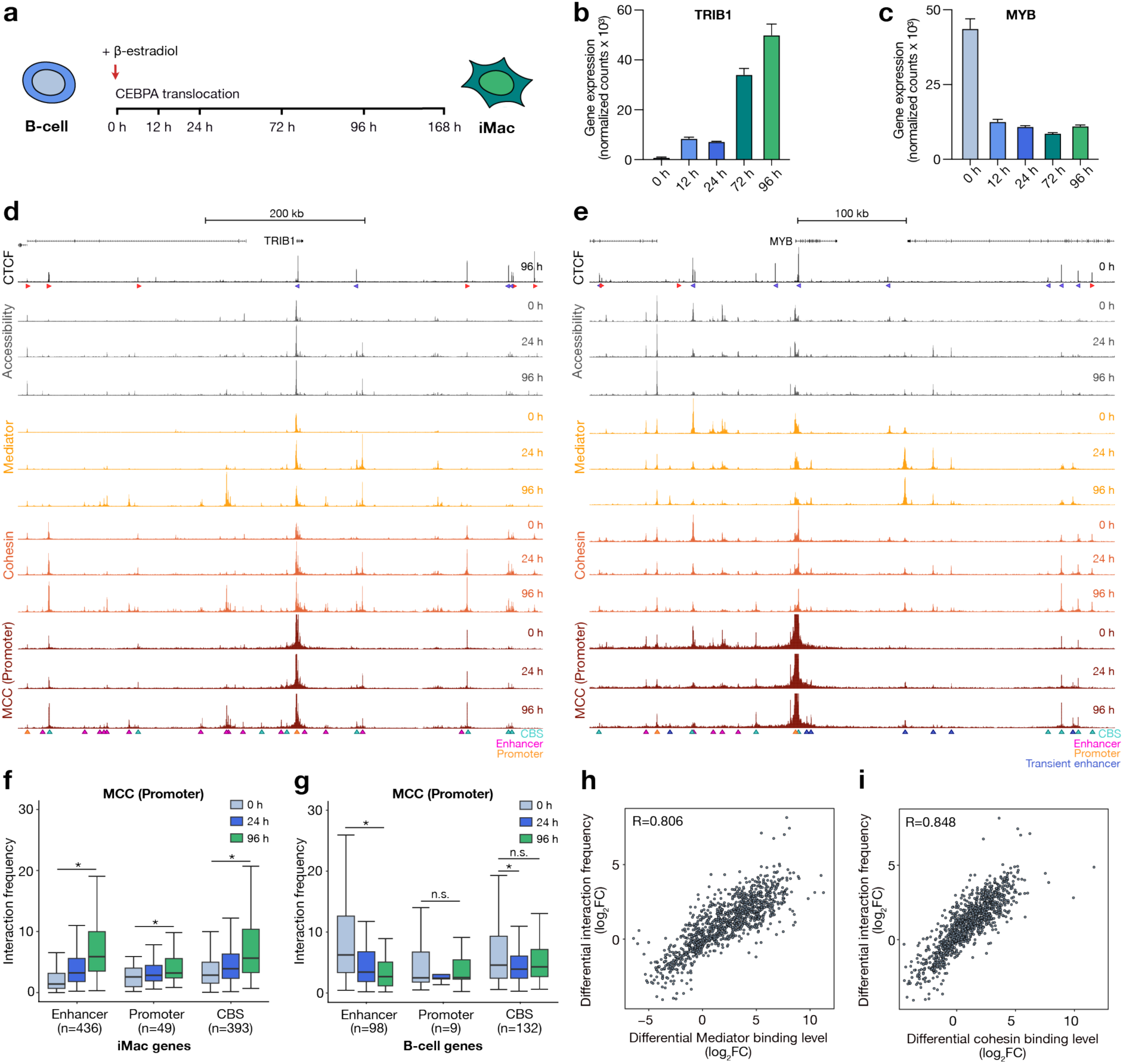
Chromatin architecture through lymphoid-to-myeloid transdifferentiation. (a) Schematic overview of the BLaER1 lymphoid-to-myeloid transdifferentiation system. (b) TRIB1 expression levels at 0 h, 12 h, 24 h, 72 h, and 96 h after differentiation induction. Expression levels are derived from TT-seq data. The bars represent the average of n = 2 replicates; the error bars indicate the standard deviation. (c) MYB expression levels through transdifferentiation, as described in panel b. (d) Chromatin landscape of the TRIB1 locus (chr8:125,079,965-125,739,965; 660 kb) during lymphoid-to-myeloid transdifferentiation. From top to bottom: gene annotation; CTCF occupancy (CTCF ChIP-seq) at 96 h; chromatin accessibility (ATAC-seq) at 0 h, 24 h, and 96 h; Mediator occupancy (MED26 ChIPmentation) at 0 h, 24 h, and 96 h; Cohesin occupancy (SMC1A ChIPmentation) at 0 h, 24 h, and 96 h; Micro-Capture-C (MCC) data from the viewpoint of the TRIB1 promoter at 0 h, 24 h, and 96 h. The axes of the profiles are scaled to signal and have the following ranges: CTCF = 0–12178; Accessibility = 0–8276; Mediator = 0–5726; Cohesin = 0–2129; MCC = 0–40. The orientations of CTCF motifs at prominent CTCF-binding sites (CBSs) are indicated by arrowheads (forward orientation in red; reverse orientation in blue). MCC interactions with CBSs, enhancers, and promoters are annotated with cyan, magenta, and orange triangles, respectively. (e) Chromatin landscape of the MYB locus (chr6:134,992,474-135,472,474; 480 kb) during lymphoid-to-myeloid transdifferentiation. Data as described in panel d, except that CTCF ChIP-seq data are shown at 0 h and that MCC interactions with transient enhancers are annotated with blue triangles. The axes of the profiles are scaled to signal and have the following ranges: CTCF = 0–6052; Accessibility = 0–3506; Mediator = 0–2277; Cohesin = 0–1247; MCC = 0–30. (f) Interaction frequencies of promoters of iMac-specific genes with enhancers, promoters, and CBSs at 0 h, 24 h, and 96 h after differentiation induction. Boxplots show the interquartile range (IQR) and median of the data; whiskers indicate the minima and maxima within 1.5 * IQR; asterisks indicate significance (P < 0.01, two-sided paired Wilcoxon signed rank test). (g) Interaction frequencies of promoters of B-cell-specific genes, as described in panel f. (h) Correlation between differential enhancer-promoter interaction frequencies and differential Mediator binding levels at the interacting elements (24 h vs 0h and 96 h vs 0 h), based on Spearman’s correlation test. (i) Correlation between differential enhancer-promoter interaction frequencies and differential cohesin binding levels at the interacting elements, as described in panel h.

Integration of these high-resolution data provides insight into the timing of regulatory events and their relationship during cellular differentiation. The *TRIB1* locus, which encodes an ubiquitin ligase adapter protein with a critical role in macrophage differentiation^56^, provides a representative example of a locus that is activated during BLaER1 transdifferentiation (**Fig. 1d and Extended Data Fig. 1c**). Our data show that the activation of *TRIB1* is associated with a gradual increase in accessibility and occupancy of H3K27ac, Mediator, and cohesin at regulatory regions, whereas CTCF binding is generally more stable. As previously described, we observe strong cohesin accumulation at CBSs^57^ and slightly weaker cohesin enrichment at Mediator-bound sites^58^. In addition, we observe changes in the structural conformation of the locus during differentiation in both the Capture-C and MCC data, which involve a gradual increase in chromatin interactions between the *TRIB1* promoter and putative enhancer elements, CBSs, and other promoters in a ∼600 kb TAD. The high-resolution of the MCC data allows for distinguishing interactions between *cis*-regulatory elements in close proximity and facilitates systematic calling of chromatin interactions^43^. Quantification of these interactions across the targeted loci shows that the described interaction pattern for the *TRIB1* locus is characteristic for upregulated loci (**Fig. 1f**).

The *MYB* oncogene locus, which encodes a B-cell-associated transcription factor^59^, provides a representative example of a locus that is de-activated during BLaER1 transdifferentiation (**Fig. 1e and Extended Data Fig. 1d**). Downregulation of *MYB* is associated with a reduction in chromatin accessibility, H3K27ac, and Mediator and cohesin binding at most putative enhancer elements in the ∼400 kb TAD. The interactions between the *MYB* promoter and these decommissioned elements gradually decrease as the cells transition into iMacs. Systematic quantification of chromatin interactions across the targeted loci shows a similar pattern for other downregulated loci (**Fig. 1g**). However, in contrast to upregulated loci, we do not observe consistent changes in promoter interactions with CBSs and other promoters in downregulated loci during differentiation (**Fig. 1g**). Interestingly, we also observe *cis-* regulatory elements that interact more frequently with the *MYB* promoter as it is de-activated. These elements are not characterized by repressive chromatin marks such as H3K27me3 (**Extended Data Fig. 1d**). Instead, they transiently gain chromatin accessibility, H3K27ac, and binding of Mediator and cohesin. We therefore speculate that these elements may function as transient enhancers that modulate the kinetics of gene silencing, as recently described in the context of erythroid differentiation^60^, although further characterization is required to confirm this. We observe elements with similar characteristics in approximately 60% of the targeted downregulated loci.

The *TRIB1* and *MYB* interaction patterns suggest that the temporal dynamics of gaining and losing Mediator and cohesin binding at enhancer elements correspond closely to the strengthening and weakening of enhancer-promoter interactions over the course of differentiation (**Fig. 1d,e**). Quantification of these patterns across all targeted loci shows a strong correlation between differential binding levels of Mediator and cohesin and enhancer-promoter interaction frequencies, signified by correlation coefficients of 0.81 and 0.85, respectively (**Fig. 1h,i**). Interestingly, we find that H3K27ac and enhancer RNA (eRNA) levels, which are more commonly used as proxies for enhancer activity, have a weaker correlation with enhancer-promoter interactions, with coefficients of 0.59 and 0.50, respectively (**Extended Data Fig. 1f,g**). This may be explained by a lower precision and dynamic range of H3K27ac as compared to Mediator and cohesin binding and technical challenges in robustly identifying eRNA levels. Together, the integration of high-resolution 3C and ChIP data allows us to characterize the 3D *cis-*regulatory landscapes of targeted tissue-specific gene loci through lymphoid-to-myeloid transdifferentiation in detail (**Extended Data Fig. 1h-k**).

### Interaction patterns across classes of *cis-*regulatory elements

Many of the gene promoters that we targeted for MCC analysis have a CBS within 5 kb. It has previously been suggested that such promoter-proximal CBSs (ppCBSs) promote long-range enhancer-promoter communication^61^ and may have a distinct function compared to CBSs at TAD boundaries (bCBSs)^62^. However, the interaction patterns of ppCBSs have not been described, as they cannot be distinguished from those of promoters by most 3C techniques. Since MCC supports base-pair resolution analysis, we leveraged this approach to compare the interaction profiles from the viewpoints of promoters and ppCBSs (if present) in the targeted loci over the course of lymphoid-to-myeloid transdifferentiation. In addition, we generated interaction profiles from the viewpoints of bCBSs in all selected loci.

The *NFKBIZ* locus, which encodes a regulator of NF-κB signaling that has a role in controlling macrophage functioning^63^, provides a representative example of an upregulated locus with distinct interaction patterns for the promoter, ppCBS, and bCBS viewpoints (**Fig. 2a,b**); the *ARL4C* locus, which is highly expressed in B-cells, provides a representative example of a downregulated locus (**Extended Data Fig. 2a,b**). In the *NFKBIZ* locus, we observe the gradual formation of relatively long-range enhancer-promoter interactions during differentiation. To investigate the relationship between the presence of a ppCBS and long-range enhancer-promoter communication, we compared the distance distribution of enhancers interacting with gene promoters with and without a ppCBS (**Fig. 2c**). This analysis confirms that promoters with a ppCBS form interactions with enhancers over larger distances compared to promoters without a ppCBS. In addition, we compared the frequencies of long-range (> 150 kb) enhancer-promoter interactions to enhancer-promoter interactions across all distances in iMac-specific loci with and without a ppCBS over the differentiation course (**Extended Data Fig. 2c**). This analysis shows that genes without a ppCBS form stronger interactions with enhancers overall compared to genes with a ppCBS. These interactions are present to some degree in B-cells and further strengthen over the differentiation course. In contrast, the loci containing ppCBSs form stronger long-range interactions, which are not pre-established and form specifically during differentiation.

**Fig. 2:**
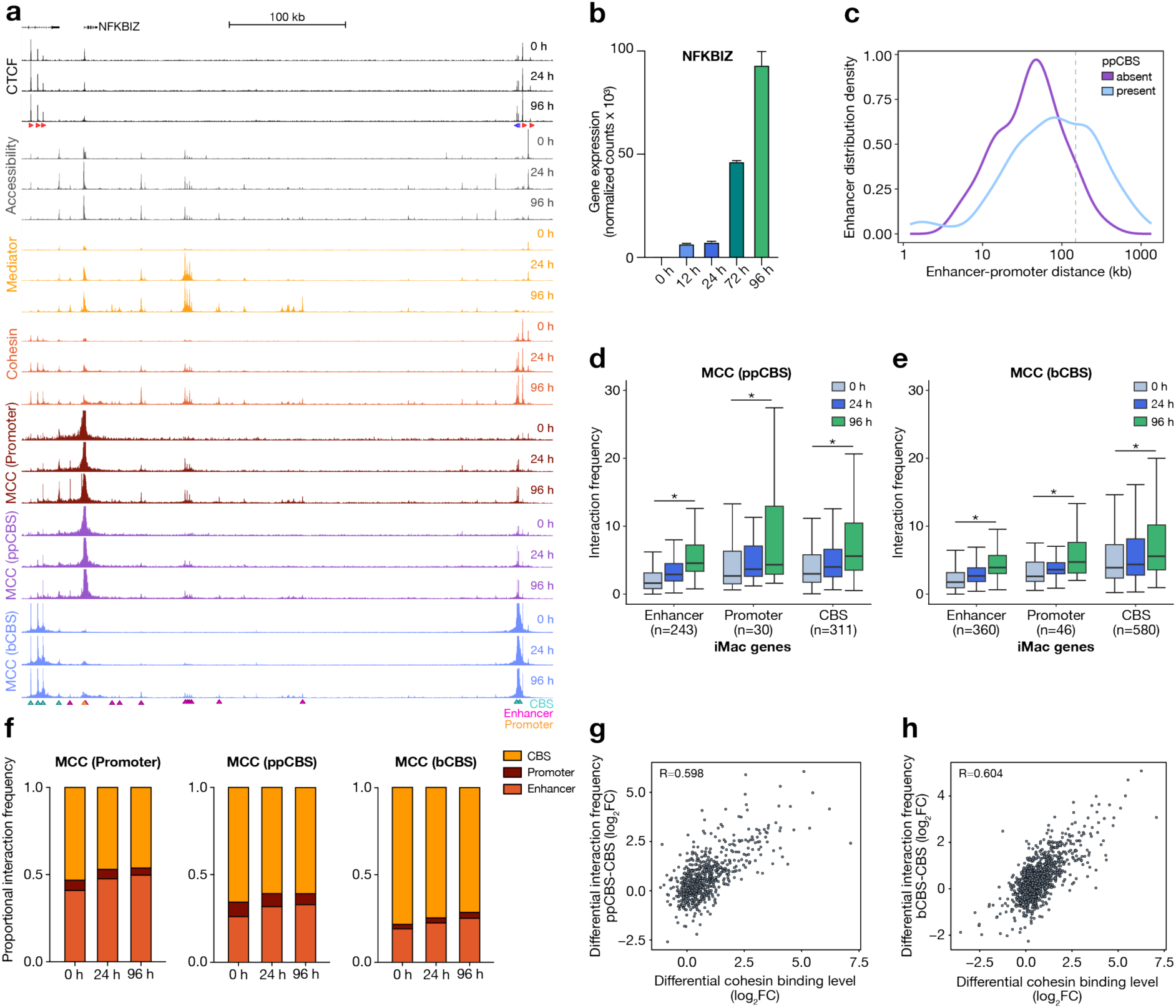
Interaction patterns across classes of cis-regulatory elements. (a) Chromatin landscape of the NFKBIZ locus (chr3:101,795,424-102,255,424; 460 kb) during lymphoid-to-myeloid transdifferentiation. From top to bottom: gene annotation; CTCF occupancy (CTCF ChIP-seq) at 0 h, 24 h, and 96 h; chromatin accessibility (ATAC-seq) at 0 h, 24 h, and 96 h; Mediator occupancy (MED26 ChIPmentation) at 0 h, 24 h, and 96 h; Cohesin occupancy (SMC1A ChIPmentation) at 0 h, 24 h, and 96 h; Micro-Capture-C (MCC) data from the viewpoint of the promoter, promoter-proximal CTCF-binding site (ppCBS), and boundary CTCF-binding site (bCBS) at 0 h, 24 h, and 96 h. The axes of the profiles are scaled to signal and have the following ranges: CTCF = 0–5547; Accessibility = 0– 4883; Mediator = 0–2252; Cohesin = 0–1773; MCC = 0–40. The orientations of CTCF motifs at prominent CBSs are indicated by arrowheads (forward orientation in red; reverse orientation in blue). MCC interactions with CBSs, enhancers, and promoters are annotated with cyan, magenta, and orange triangles, respectively. (b) NFKBIZ expression at 0 h, 12 h, 24 h, 72 h, and 96 h after differentiation induction. Expression levels are derived from TT-seq data. The bars represent the average of n = 2 replicates; the error bars indicate the standard deviation. (c) Distribution of enhancer-promoter distances of promoters with and without a ppCBS (± 5 kb from the promoter). The grey line marks the 150 kb threshold used to classify distal enhancers in Extended Data Fig. 2c. (d) Interaction frequencies of ppCBSs of iMac-specific genes with enhancers, promoters, and CBSs at 0 h, 24 h, and 96 h after differentiation induction. Boxplots show the interquartile range (IQR) and median of the data; whiskers indicate the minima and maxima within 1.5 * IQR; asterisks indicate significance (P < 0.01, two-sided paired Wilcoxon signed rank test). (e) Interaction frequencies of bCBSs of iMac-specific genes, as described in panel d. (f) Comparison of the proportion of interactions of promoters, ppCBSs, and bCBSs with enhancers, promoters, and CBSs in iMac-specific gene loci at 0 h, 24 h, and 96 h. (g) Correlation between differential interaction frequencies between ppCBS and CBSs and differential cohesin binding levels at the interacting elements (24 h vs 0h and 96 h vs 0 h), based on Spearman’s correlation test. (h) Correlation between differential interaction frequencies between bCBS and CBSs and differential cohesin binding levels at the interacting elements, as described in panel g.

The interaction profiles from the ppCBS viewpoint in the *NFKBIZ* locus show that the ppCBS forms interactions with the enhancers in the region, although these are weaker compared to the promoter viewpoint, and with other CBSs in the region, which are stronger compared to the promoter viewpoint. The bCBS viewpoint most strongly interacts with the CBSs at the other TAD boundary, but also forms weak interactions with the enhancers and promoters in the region. These interactions all gradually increase over the differentiation course. Systematic quantification of chromatin interactions in the targeted loci shows a similar pattern across the upregulated loci (**Fig. 2d,e**) and the reverse in the downregulated loci (**Extended Data Fig. 2d,e**). Comparison of these data to the interaction patterns of gene promoters (**Fig. 1f,g**) shows that promoter interactions are generally more dynamic over the differentiation course compared to ppCBS and especially bCBS interactions. For a more direct comparison of interaction patterns across the classes of *cis*-regulatory elements, we calculated changes in the relative proportions of interactions over the differentiation course (**Fig. 2f and Extended Data Fig. 2f**). This comparison shows that enhancers interact more frequently with ppCBSs than with bCBSs, which suggests that ppCBSs may have a direct role in bringing enhancers in the vicinity of gene promoters. In addition, these analyses indicate that promoters, ppCBSs, and bCBSs interact promiscuously with active *cis*-regulatory elements within TADs, although they do have distinct preferences for specific types of elements. The common feature at these *cis-*regulatory elements is enrichment of cohesin, suggesting a role for loop extrusion in mediating these interactions. In line with this hypothesis, we find that differential cohesin binding correlates well with the interaction frequencies between CBSs (**Fig. 2g,h**), with coefficients of 0.6. These correlations are lower compared to enhancer-promoter interactions (**Fig. 1i**), which may be explained by the larger dynamic range in interactions between enhancers and promoters through differentiation compared to CBSs. Of note, we find that CBS interaction frequencies correlate better with cohesin binding than with CTCF binding itself (**Extended Data Fig. 2g,h**), which is in agreement with the more constitutive binding patterns of CTCF.

### Dynamic chromatin hub formation and dissolution during differentiation

The MCC data show complex interaction patterns involving multiple promoters, enhancers, and CBSs in the targeted regions. However, since MCC is based on the detection of pair-wise interactions, these data cannot resolve how these elements interact together in higher-order 3D chromatin structures. To analyze 3D interactions between *cis*-regulatory elements, we used the Tri-C technique to measure multi-way interactions in the 51 targeted regions of interest. Tri-C uses the restriction enzyme *Nla*III for chromatin digestion and therefore generates lower-resolution data compared to MCC, which complicates direct quantitative comparisons between these datasets. Because Tri-C requires the restriction fragments at the targeted viewpoints to be relatively small (< 300 bp) to facilitate efficient detection of multiple interactions^44^, we targeted enhancers instead of promoters to assess multi-way enhancer-promoter interactions, as this provides more flexibility to select suitable viewpoints. In addition to enhancers, we targeted bCBSs in the selected regions. The Tri-C interactions are represented in viewpoint-specific contact matrices, in which the frequencies with which two chromatin fragments interact simultaneously with the viewpoint are plotted (**Fig. 3a**). Mutually exclusive interactions are depleted from these matrices, whereas preferential simultaneous interactions between *cis-*regulatory elements in higher-order structures are visible as enrichments at the intersections between these elements. Note that these matrices typically show strong signals along the proximity-excluded region at the viewpoint, which represent viewpoint-proximal interactions throughout the targeted region. Since regions in close genomic proximity are expected to form strong interactions, these stripe patterns do not represent specific higher-order chromatin conformations.

**Fig. 3:**
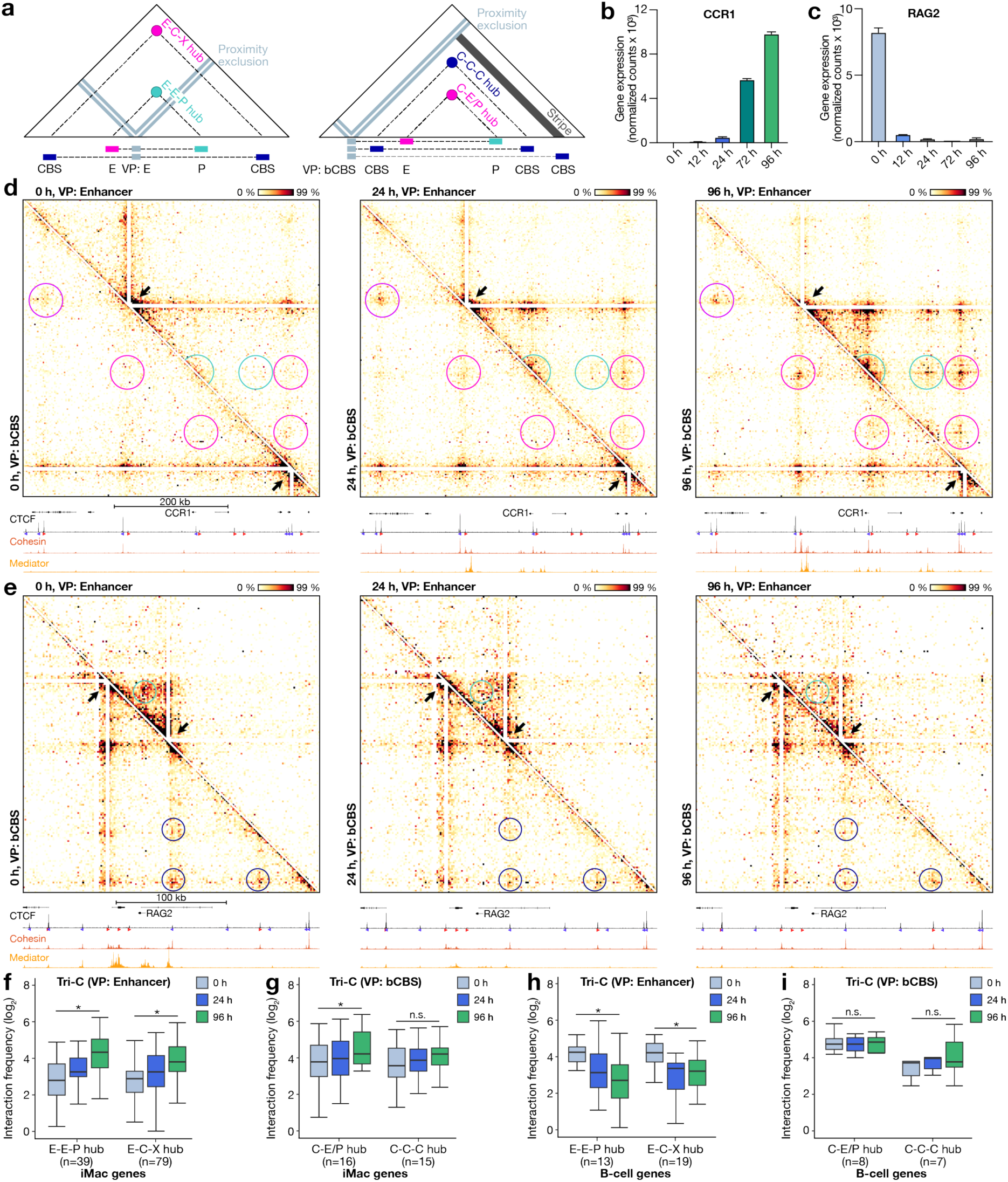
Dynamic chromatin hub formation and dissolution during differentiation. (a) Schematic overview of Tri-C data visualization. Viewpoint-specific contact matrices show the frequency with which two regions interact simultaneously with the viewpoint. The proximity signal around the viewpoint is excluded. Quantified regions for the enhancer viewpoints include three-way interactions involving two enhancers and a promoter (E-E-P hubs) and three-way interactions involving an enhancer, CTCF-binding site (CBS), and any other cis-regulatory element (E-C-X hubs). Quantified regions for the CBS viewpoints include three-way interactions involving three CBSs (C-C-C hubs) and three-way interactions involving a CBS and any combination of enhancers and/or promoters (C-E/P hubs). (b) CCR1 expression at 0 h, 12 h, 24 h, 72 h, and 96 h after differentiation induction. Expression levels are derived from TT-seq data. The bars represent the average of n = 2 replicates; the error bars indicate the standard deviation. (c) RAG2 expression levels through transdifferentiation, as described in panel b. (d) Tri-C contact matrices of the CCR1 locus (chr3:45,902,299-46,427,299; 525 kb; 2.5 kb resolution) during lymphoid-to-myeloid transdifferentiation at 0 h, 24 h, and 96 h. The top-right matrix shows Tri-C data from the viewpoint of an enhancer; the bottom-left matrix shows the viewpoint of a boundary CTCF-binding site (bCBS). The viewpoints are indicated with white triangles. E-E-P, E-C-X, and C-E/P contacts are highlighted in cyan, magenta, and dark blue circles, respectively. The profiles below the Tri-C contact matrices show occupancy of CTCF (CTCF ChIP-seq), Mediator (MED26 ChIPmentation), and cohesin (SMC1A ChIPmentation) at corresponding time points. The axes are scaled to signal and have the following ranges: CTCF = 0–10069; Cohesin = 0–2960; Mediator = 0–6075. (e) Tri-C contact matrices of the RAG2 locus (chr11:36,486,822-36,756,822; 170 kb; 1.5 kb resolution) during lymphoid-to-myeloid transdifferentiation, as described in panel d. The axes of the profiles are scaled to signal and have the following ranges: CTCF = 0–14993; Cohesin = 0–2868; Mediator = 0–2831. (f) Multi-way interaction frequencies of E-E-P and E-C-X hubs in iMac-specific loci at 0 h, 24 h, and 96 h after differentiation induction. Boxplots show the interquartile range (IQR) and median of the data; whiskers indicate the minima and maxima within 1.5 * IQR; asterisks indicate significance (P < 0.01, two-sided paired Wilcoxon signed rank test). (g) Multi-way interaction frequencies of C-E/P and C-C-C hubs in iMac-specific loci, as described in panel f. (h) Multi-way interaction frequencies of E-E-P and E-C-X hubs in B-cell-specific loci, as described in panel f. (i) Multi-way interaction frequencies of C-E/P and C-C-C hubs in B-cell-specific loci, as described in panel f.

The *CCR1* locus, which encodes a chemokine receptor that is highly expressed in macrophages^64^, and the *RAG2* locus, which encodes an enzyme involved in VDJ recombination in B-cells^65^, provide representative examples of the establishment and dissolution of higher-order chromatin structures through lymphoid-to-myeloid transdifferentiation (**Fig. 3b-e**). At the *CCR1* locus, we observe the formation of a hub structure, characterized by simultaneous interactions between the enhancer viewpoint, the *CCR1* promoter, and other enhancer clusters in the region (**Fig. 3d, top matrix, cyan circles**). Interestingly, we observe that the CBSs in the region are also included in these hubs (**Fig. 3d, top matrix, magenta circles**). The contact matrices from the viewpoint of the CBS at the downstream boundary of the *CCR1* TAD confirm the cooperative interactions between the promoter, enhancers, and CBSs in chromatin hubs in the *CCR1* locus (**Fig. 3d, bottom matrix, magenta circles**). Both the enhancer and the CBS viewpoint in the *CCR1* locus show that these chromatin hubs form gradually over the differentiation course. Tri-C data in the *NFKBIZ* and *TRIB1* loci (**Extended Data Fig. 3a,b**) and quantification of the multi-way chromatin interactions across all upregulated loci (**Fig. 3f,g**) show a similar pattern of gradual chromatin hub formation. Analysis of the CBS contact matrices across the targeted loci shows that these are characterized by distinct interaction foci as well as stripes that emanate from CBSs throughout the region and likely represent active loop extrusion by cohesin molecules (**Fig. 3d,e and Extended Data Fig. 3a-d**). We observe that loci with relatively little CTCF binding, such as *NFKBIZ*, are characterized by CBS stripes, whereas CTCF-dense regions, such as *CCR1*, also form clear CBS foci. The *RAG2* locus shows that cooperative interactions between enhancers and promoters gradually dissolve as *RAG2* is downregulated (**Fig. 3e, top matrix, cyan circles**). In contrast, multi-way CBS interactions are relatively stable over the differentiation course (**Fig. 3e, bottom matrix, dark blue circles**), which is in line with the relatively stable pair-wise CBS interactions identified by MCC (**Fig. 1e,g**). Quantification of the multi-way chromatin interactions shows a similar pattern across the targeted downregulated loci (**Fig. 3h,i**). We detect multi-way chromatin interactions with enhancers and CBSs in the majority of the targeted up- and downregulated loci (**Extended Data Fig. 3e-f**). Together, the Tri-C data therefore show that the clustering of *cis-*regulatory elements into higher-order chromatin hubs is a common feature of tissue-specific gene loci, and that these hubs form and dissolve dynamically as genes are up- and downregulated during cellular differentiation.

### CTCF supports the formation of chromatin hubs but is not required for pair-wise enhancer-promoter interactions

The incorporation of CBSs into chromatin hubs suggests that CTCF may have a function in the formation of these hubs and, more generally, in the regulation of enhancer-promoter communication during cellular differentiation. To test this, we performed 3C experiments in iMacs after auxin-mediated CTCF depletion^54^ throughout the differentiation course (96 hours; **Fig. 4a**). To confirm efficient depletion, we mapped CTCF binding in auxin- and control-treated iMacs (**Extended Data Fig. 4a**). In addition, we confirmed that the BLaER1 cells are still efficiently converted into iMacs in absence of CTCF by measuring specific B-cell and macrophage marker genes with qPCR through the differentiation course (**Extended Data Fig. 4b**).

**Fig. 4:**
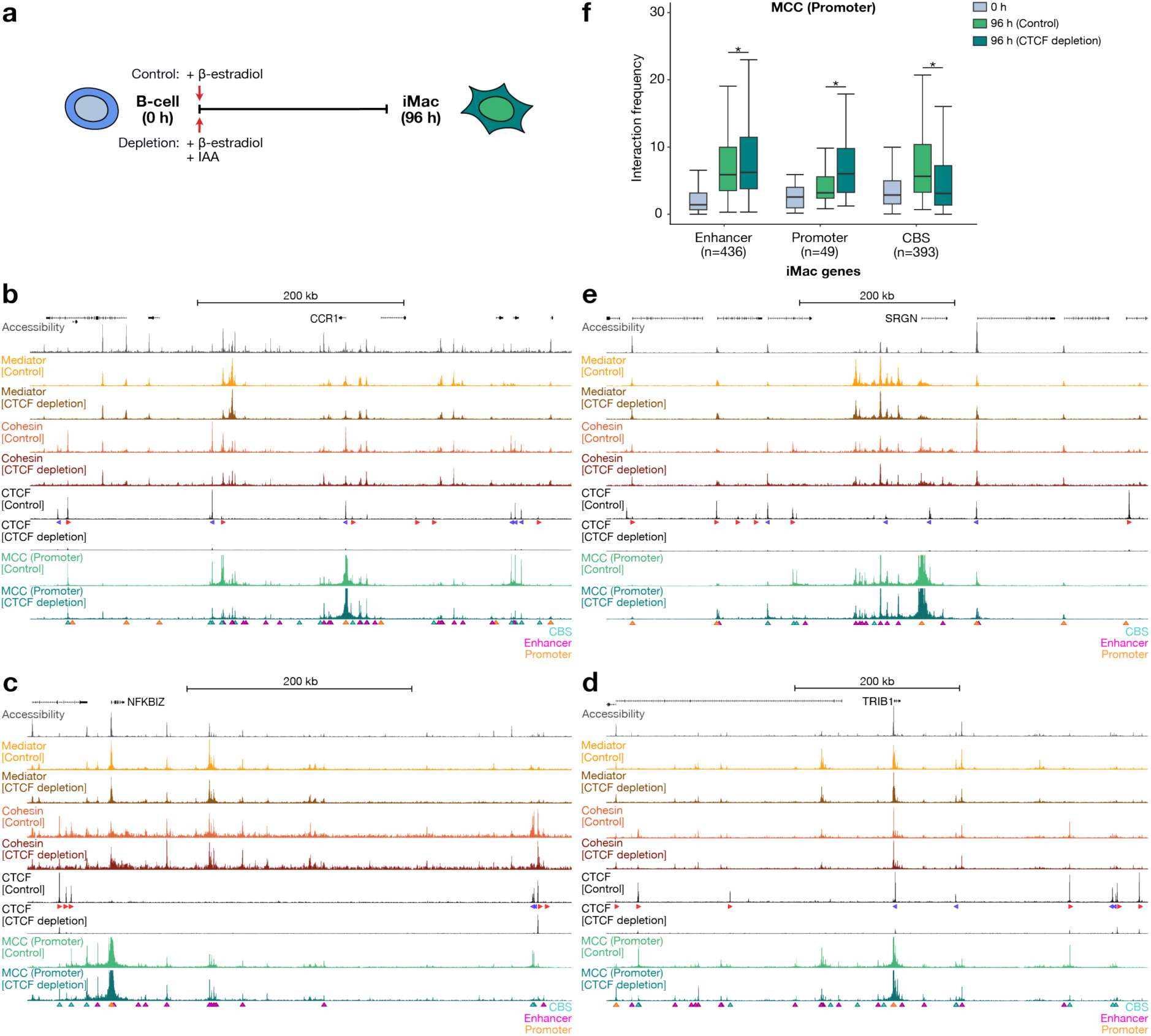
CTCF is not required for pair-wise enhancer-promoter interactions. (a) Schematic overview of CTCF depletion during lymphoid-to-myeloid transdifferentiation. (b) Chromatin interactions in the CCR1 locus (chr3:45,902,299-46,427,299; 525 kb) in control and CTCF-depleted cells at 96 h after differentiation induction. From top to bottom: gene annotation; chromatin accessibility (ATAC-seq); Mediator occupancy (MED26 ChIPmentation); Cohesin occupancy (SMC1A ChIPmentation); CTCF occupancy (CTCF ChIPmentation); Micro-Capture-C (MCC) data from the viewpoint of the promoter. The axes of the profiles are scaled to signal and have the following ranges: Accessibility = 0–1565; Mediator = 0–1463; Cohesin = 0–975; CTCF = 0–468; MCC = 0–40. The orientations of CTCF motifs at prominent CBSs are indicated by arrowheads (forward orientation in red; reverse orientation in blue). MCC interactions with CBSs, enhancers, and promoters are annotated with cyan, magenta, and orange triangles, respectively. (c) Chromatin interactions in the NFKBIZ locus (chr3:101,766,932-102,266,932; 500 kb) in control and CTCF-depleted cells, as described in panel b. The axes of the profiles are scaled to signal and have the following ranges: Accessibility = 0–4883; Mediator = 0–760; Cohesin = 0–367; CTCF = 0–286; MCC = 0–40. (d) Chromatin interactions in the TRIB1 locus (chr8:125,079,965-125,739,965; 660 kb) in control and CTCF-depleted cells, as described in panel b. The axes of the profiles are scaled to signal and have the following ranges: Accessibility = 0–8276; Mediator = 0–1858; Cohesin = 0–1660; CTCF = 0–468; MCC = 0–40. (e) Chromatin interactions in the SRGN locus (chr10:68,884,514-69,234,514; 350 kb) in control and CTCF-depleted cells, as described in panel b. The axes of the profiles are scaled to signal and have the following ranges: Accessibility = 0– 5900; Mediator = 0–1430; Cohesin = 0–777; CTCF = 0–318; MCC = 0–50. (f) Interaction frequencies of promoters of iMac-specific genes with enhancers, promoters, and CBSs at 0 h and at 96 h in control and CTCF-depleted cells. Boxplots show the interquartile range (IQR) and median of the data; whiskers indicate the minima and maxima within 1.5 * IQR; asterisks indicate significance (P < 0.01, two-sided paired Wilcoxon signed rank test).

To assess the effects of CTCF depletion on the gradual formation of enhancer-promoter interactions during lymphoid-to-myeloid transdifferentiation, we performed MCC, with the gene promoters in the selected loci as viewpoints. Analysis of the *CCR1*, *NFKBIZ*, and *TRIB1* loci shows that, as expected, CTCF depletion leads to a strong reduction in interactions between promoters and CBSs (**Fig. 4b-d**). In contrast, the interactions between promoters and enhancers in these loci are not strongly affected. As exemplified in the *TRIB1* and *SRGN* loci, we observe that many promoter-promoter interactions are increased after CTCF depletion, both within and beyond TAD boundaries (**Fig. 4d,e**). Systematic quantification of the MCC interactions shows similar patterns as in the described examples (**Fig. 4f**). Across upregulated loci, promoter-CBS interactions are weakened, enhancer-promoter interactions are relatively stable, and promoter-promoter interactions are increased after CTCF depletion.

Given the strong correlation between differential Mediator and cohesin binding levels and enhancer-promoter interaction frequencies during differentiation, we performed additional ChIPmentation experiments after CTCF depletion to investigate whether we can explain the observed changes in interaction patterns by changes in the binding levels of Mediator and cohesin. As expected, CTCF depletion leads to a reduction in cohesin occupancy at CBSs (**Fig. 4b-e**). In contrast, we observe minor changes in the distribution of Mediator and cohesin at enhancers, which do not correlate well with the minor changes we observe in enhancer-promoter interactions after CTCF depletion (**Extended Data Fig. 4c,d**).

To characterize the role of CTCF in the formation of chromatin hubs during cellular differentiation, we performed Tri-C analysis, with the enhancers in the selected loci as viewpoints. In stark contrast to the minor effects of CTCF depletion on enhancer-promoter interactions measured by MCC, the Tri-C contact matrices of the *CCR1*, *NFKBIZ*, and *TRIB1* loci show drastic changes in higher-order 3D chromatin structures following CTCF depletion (**Fig. 5a-c**). In absence of CTCF, the formation of cooperative interactions between the targeted enhancers and other *cis*-regulatory elements in these regions is impaired. Unsurprisingly, cooperative interactions that involve a CBS are most strongly reduced. However, interestingly, cooperative interactions involving only enhancers and promoters are also decreased. This effect appears stronger in CBS-dense regions (e.g., *CCR1*) compared to regions with relatively little CTCF binding (e.g., *NFKBIZ*), although it is detectable across all targeted regions. It is important to note that the loss of these interactions is unlikely to reflect changes in the distribution of the pair-wise interactions with the targeted enhancer viewpoint, since the MCC data show that enhancer-promoter interactions remain relatively stable (and in some cases even increase) after CTCF depletion (**Fig. 4f**).

**Fig. 5:**
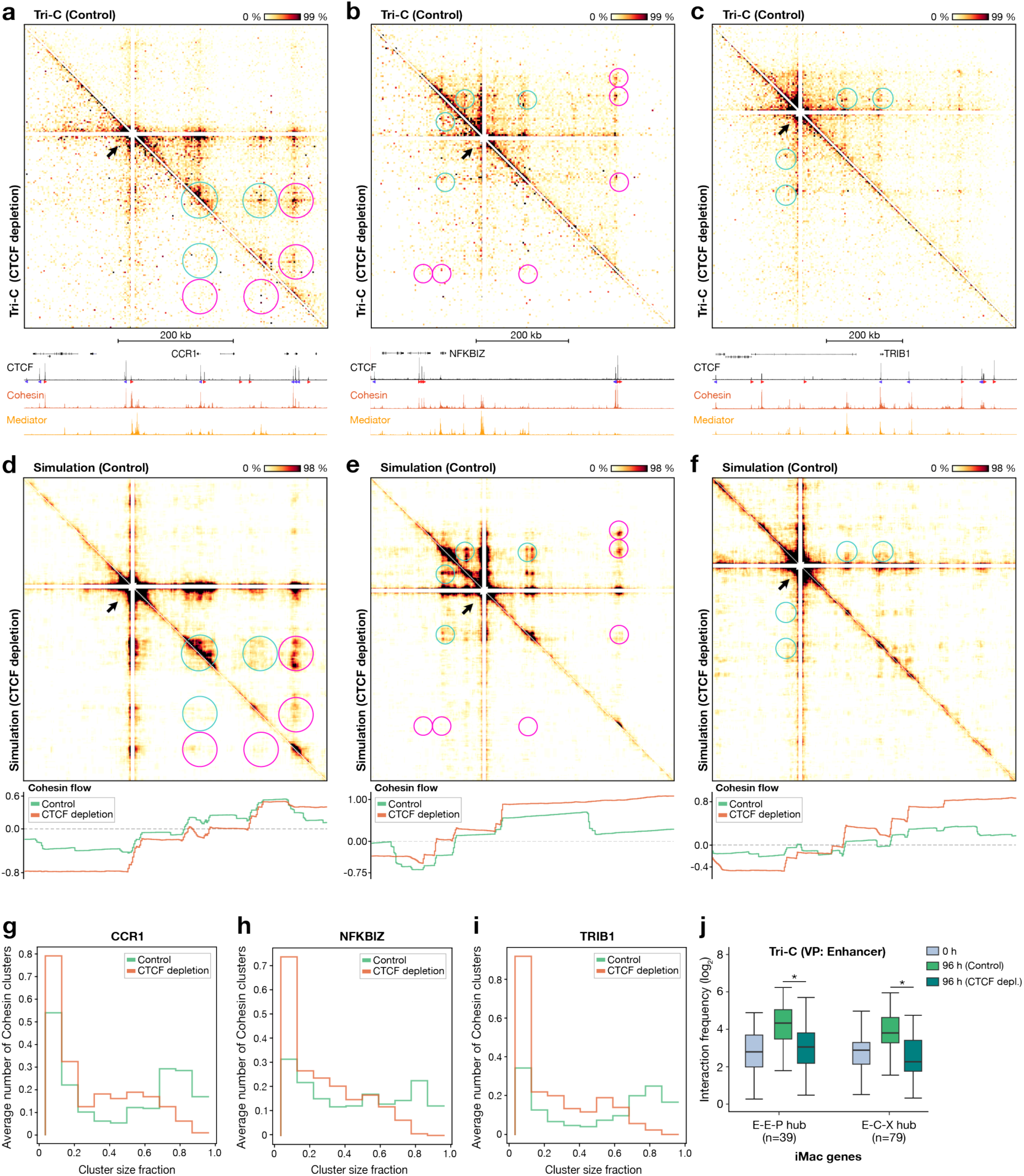
CTCF supports the formation of chromatin hubs. (a) Tri-C contact matrices of the CCR1 locus (chr3:45,902,299-46,427,299; 525 kb; 2.5 kb resolution) from the viewpoint of an enhancer in control (top-right matrix) and CTCF-depleted (bottom-left matrix) cells at 96 h after differentiation induction. The viewpoints are indicated with white triangles. E-E-P and E-C-X contacts are highlighted in cyan and magenta circles, respectively. The profiles below the Tri-C contact matrices show occupancy of CTCF (CTCF ChIP-seq), Mediator (MED26 ChIPmentation), and cohesin (SMC1A ChIPmentation) at 96 h. The axes of the profiles are scaled to signal and have the following ranges: CTCF = 0–10069; Cohesin = 0–2960; Mediator = 0–6075. (b) Tri-C contact matrices of the NFKBIZ locus (chr3:101,699,405-102,350,405; 651 kb; 3.5 kb resolution) in control and CTCF-depleted cells at 96 h, as described in panel a. The axes of the profiles are scaled to signal and have the following ranges: CTCF = 0–5547; Cohesin = 0–1773; Mediator = 0–2252. (c) Tri-C contact matrices of the TRIB1 locus (chr8:124,988,732-125,788,732; 800 kb; 4 kb resolution) in control and CTCF-depleted cells at 96 h, as described in panel a. The axes are scaled to signal and have the following ranges: CTCF = 0–12178; Cohesin = 0–2129; Mediator = 0–5726. (d) Tri-C contact matrices generated with molecular dynamics simulations of the CCR1 locus (genomic coordinates as described in panel a; 2 kb resolution) in control (top-right matrix) and CTCF-depleted (bottom-left matrix) cells at 96 h after differentiation induction. The profile below shows cohesin flow (net number of cohesin molecules moving through the locus per minute) in control and CTCF-depleted conditions. Positive values indicate cohesin movement in sense direction; negative values indicate antisense direction; extreme values imply less constrained cohesin movement. (e) Tri-C contact matrices generated with molecular dynamics simulations of the NFKBIZ locus (genomic coordinates as described in panel b; 2 kb resolution) in control and CTCF-depleted cells at 96 h and cohesin flow as described in panel d. (f) Tri-C contact matrices generated with molecular dynamics simulations of the TRIB1 locus (genomic coordinates as described in panel c; 2 kb resolution) and cohesin flow as described in panel d. (g) Cohesin cluster size frequencies, measured as fraction of the total number of molecules from the simulations of the CCR1 locus. (h) Cohesin cluster size frequencies in the NFKBIZ locus, as described in panel g. (i) Cohesin cluster size frequencies in the TRIB1 locus, as described in panel g. (j) Frequencies of three-way interactions involving two enhancers and a promoter (E-E-P hub) and three-way interactions involving an enhancer, CTCF-binding site, and any other cis-regulatory element (E-C-X hub) in iMac-specific loci at 0 h and at 96 h in control and CTCF-depleted cells. Boxplots show the interquartile range (IQR) and median of the data; whiskers indicate the minima and maxima within 1.5 * IQR; asterisks indicate significance (P < 0.01, two-sided paired Wilcoxon signed rank test).

We used molecular dynamics simulations of 3D chromatin folding in the *CCR1*, *NFKBIZ*, and *TRIB1* loci to obtain a more comprehensive understanding of the effects of CTCF depletion. To this end, we built on our existing modeling framework^66,67^ to model a 2 Mb region around each of these three genes (**Methods**). From these simulations, we extracted multi-way interactions with the same enhancers that were used as Tri-C viewpoints and produced contact matrices that closely resemble the experimental Tri-C results (**Fig. 5a-f and Extended Data Fig. 5a-c**). In presence of CTCF, the targeted enhancers engage in multi-way interactions involving both their cognate gene promoter and other enhancers, as well as CBSs. Removal of CTCF from the models does not significantly affect extracted pair-wise interactions (**Extended Data Fig. 5d,e**), yet fully reproduces the impairment of chromatin hub formation across these loci (**Fig. 5a-f and Extended Data Fig. 5f,g**). The models therefore allow us to investigate the underlying mechanisms by interrogating the distribution and net flow of extruding cohesin molecules in the three loci. Consistent with the cohesin ChIPmentation data (**Fig. 1d,e,i**), the simulations show that cohesin accumulates strongly at CBSs and weakly at active enhancers and promoters in control conditions (**Fig. 5d-f, bottom panels, green tracks**). In absence of strong CTCF-bound insulation sites, cohesin flows more freely (**Fig. 5d-f, bottom panels, orange tracks**), which results in a marked reduction of cohesin clustering to essentially only pair-wise interactions (**Fig. 5g-i**). These analyses therefore suggest that in absence of relatively long-lived CTCF residence at CBSs, the spatiotemporal clustering of cohesin changes such that the formation of higher-order chromatin hubs is no longer supported. Importantly, the other upregulated loci show similar patterns as the *CCR1*, *NFKBIZ*, and *TRIB1* loci. Cooperative interactions in CTCF-depleted iMacs are strongly reduced, with interactions directly involving CBSs (or elements in close vicinity to CBSs) most strongly affected (**Fig. 5j and Extended Data Fig. 5h-m**). Together, these results indicate that CTCF-mediated interactions provide a scaffold for the formation of chromatin hubs during cellular differentiation.

### Gene expression patterns are instructed by pair-wise enhancer-promoter interactions but not dependent on chromatin hubs

To investigate the function of CTCF-dependent chromatin structures in gene regulation, we measured the effects of CTCF depletion during lymphoid-to-myeloid transdifferentiation on gene expression. In agreement with previous reports^54,68,69^ and with the observation that BLaER1 cells still efficiently transdifferentiate into iMacs in absence of CTCF (**Extended Data Fig. 4b**), we find that CTCF depletion has modest effects on gene expression during lymphoid-to-myeloid transdifferentiation (**Fig. 6a**). Upon CTCF depletion, 718 genes are significantly downregulated and 558 genes are significantly upregulated, with a log_2_ fold-change < 2 for 90% of the significantly differentially expressed genes (**Extended Data Fig. 6a and Supplementary Table 2**). Differentially expressed genes are enriched for B-cell- and iMac-specific genes and are more likely to have a CBS near their promoter and to be regulated by distal enhancers compared to unaffected genes (**Extended Data Fig. 6b-e**). However, interestingly, we do not generally observe a significant decrease in gene expression in the upregulated gene loci in which we observe a strong impairment in chromatin hub formation, including *CCR1*, *NFKBIZ* and *TRIB1* (**Extended Data Fig. 6a**). This suggests that CTCF-dependent chromatin hubs do not have a critical role in the regulation of gene expression during cellular differentiation.

**Fig. 6:**
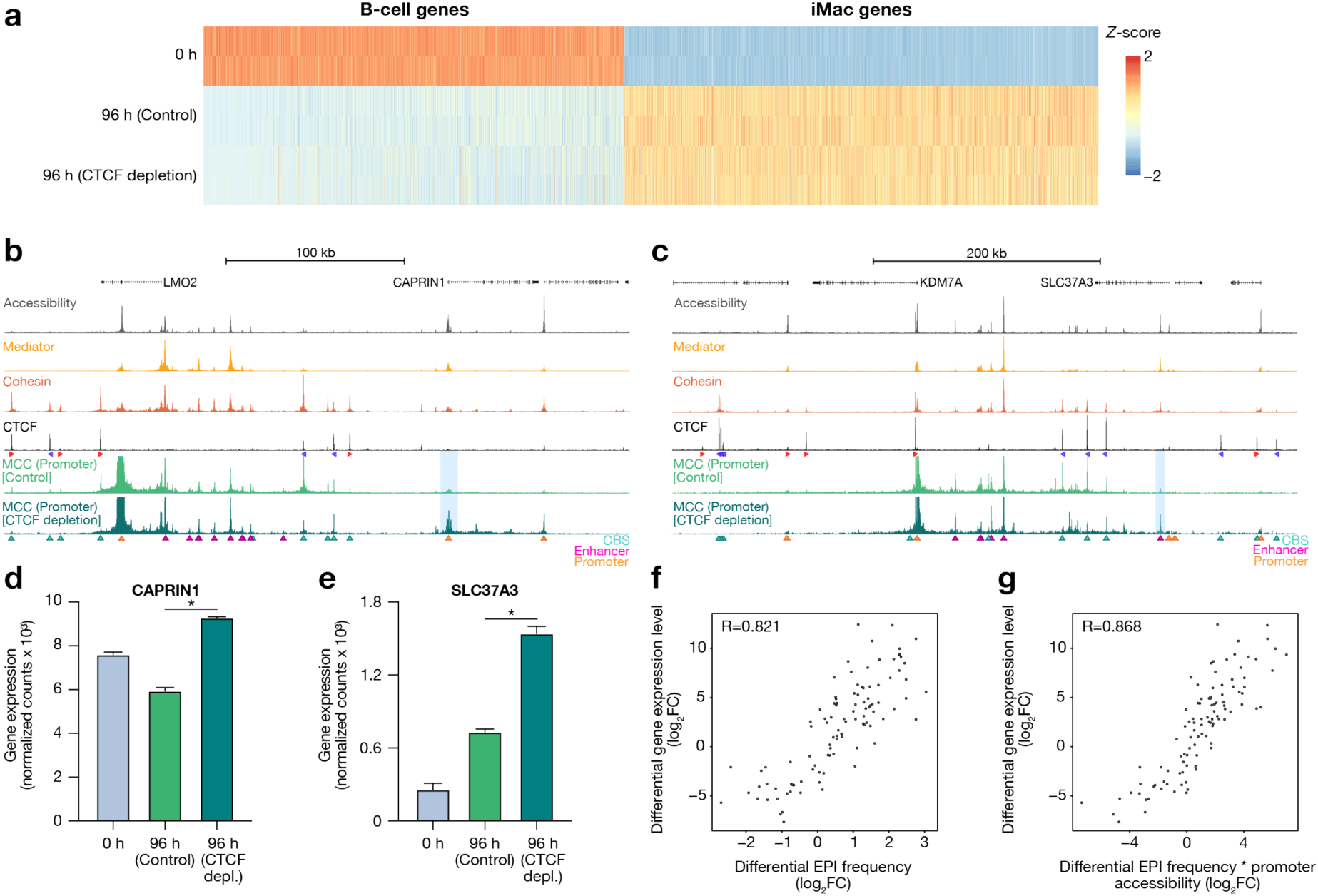
Pair-wise enhancer-promoter interactions instruct gene expression patterns. (a) Heatmap showing the Z-score of normalized RNA counts of B-cell-specific genes (n = 2968) and iMac-specific genes (n = 3341) at 0 h and at 96 h in control-treated and CTCF-depleted cells. RNA-seq experiments were performed in n = 2 replicates. (b) Chromatin interactions in the LMO2 locus (chr11:33,804,269-34,154,269; 350 kb) in control and CTCF-depleted cells at 96 h after differentiation induction. From top to bottom: gene annotation; chromatin accessibility (ATAC-seq); Mediator occupancy (MED26 ChIPmentation); cohesin occupancy (SMC1A ChIPmentation); CTCF occupancy (CTCF ChIPmentation); Micro-Capture-C (MCC) data from the viewpoint of the promoter. The axes of the profiles are scaled to signal and have the following ranges: Accessibility = 0–4018; Mediator = 0–7353; cohesin = 0–1875; CTCF = 0–12417; MCC = 0–40. The orientations of CTCF motifs at prominent CBSs are indicated by arrowheads (forward orientation in red; reverse orientation in blue). MCC interactions with CBSs, enhancers, and promoters are annotated with cyan, magenta, and orange triangles, respectively, and ectopic interactions in CTCF-depleted cells are highlighted in light blue. (c) Chromatin interactions in the KDM7A locus (chr7:139,961,429-140,511,429; 550 kb) in control and CTCF-depleted cells, as described in panel b. The axes of the profiles are scaled to signal and have the following ranges: Accessibility = 0–3566; Mediator = 0–7621; Cohesin = 0–2469; CTCF = 0–7069; MCC = 0–40. (d) CAPRIN1 expression levels at 0 h and at 96 h in control-treated and CTCF-depleted cells. Expression levels are derived from RNA-seq data. The bars represent the average of n = 2 replicates; the error bars indicate the standard deviation. Asterisks indicate significant differences (adjusted P-value < 0.01 and log2 FC > 0.6). (e) SLC37A3 expression levels at 0 h and at 96 h in control-treated and CTCF-depleted cells, as described in panel d. (f) Correlation between differential expression levels and total enhancer-promoter interaction frequencies (24 h vs 0h and 96 h vs 0 h), based on Spearman’s correlation test. (g) Correlation between differential expression levels and the product of total enhancer-promoter interaction frequencies and promoter accessibility levels, as described in panel f.

It has previously been shown that CBSs contribute to the specificity of enhancer-promoter communication by preventing interactions between enhancers and promoters across TAD borders^70^. In agreement with this model, we observe that CTCF depletion leads to the formation of ectopic interactions in two of the targeted upregulated loci. In the *LMO2* locus, we observe increased interactions with the promoter of *CAPRIN1* (**Fig. 6b**); in the *KDM7A* locus, we observe increased interactions with *cis-*regulatory elements of the *SLC37A3* gene (**Fig. 6c**). These rewired interactions are associated with a significant increase in *CAPRIN1* and *SLC37A3* expression (**Fig. 6d,e**). In addition to the ectopic interactions in the *LMO2* and *KDM7A* loci, we find subtle changes in interaction profiles in some of the other targeted loci, which are also associated with small changes in gene expression. For example, in the *IRF8* and *MAFB* locus, both enhancer-promoter interactions and gene expression levels are slightly decreased and increased, respectively, after CTCF depletion (**Extended Data Fig. 6a,f,g**). Furthermore, consistent with the observation that downregulated genes are more likely to have distal enhancers (**Extended Data Fig. 6d**), we find that decreased enhancer-promoter interactions upon CTCF depletion are more likely to be distal (> 150 kb) from the promoter compared to increased enhancer-promoter interactions (**Extended Data Fig. 6e**).

It is of interest that both gene expression patterns and pair-wise enhancer-promoter interactions are relatively stable after CTCF depletion and that the minor changes in gene expression following CTCF depletion can often be explained by changes in pair-wise enhancer-promoter interaction frequencies. This suggests that basic contacts between enhancers and promoters (in the form of pair-wise interactions) are more relevant for gene regulation than their cooperative interactions within chromatin hubs. To further explore the relationship between pair-wise enhancer-promoter interactions and gene expression, we computed a total enhancer-promoter interaction score for the targeted gene promoters based on the MCC data through lymphoid-to-myeloid transdifferentiation. Interestingly, we find that changes in enhancer-promoter interactions correlate very well with changes in gene expression over the differentiation course, with a coefficient of 0.82 (**Fig. 6f**). Integration of the accessibility of the gene promoters further increases this correlation, to a coefficient of 0.87 (**Fig. 6g**). Together, these results indicate that the gradual formation and dissolution of enhancer-promoter interactions are predictive of dynamic changes in gene expression during cellular differentiation, but that these gene expression changes do not strongly depend on the higher-order configuration of enhancers and promoters in CTCF-dependent hub structures.

## DISCUSSION

In this study, we characterize structural and functional features of the genome over the course of lymphoid-to-myeloid transdifferentiation. We find that interactions between promoters, enhancers, and CBSs gradually form and dissolve in up- and downregulated gene loci during differentiation. For enhancer-promoter interactions, we find that their frequencies correlate very well with changes in the levels of Mediator and cohesin binding at these elements. This is consistent with previous studies that have suggested a role for Mediator^71,72^ and cohesin-mediated loop extrusion^73–76^ in the formation of enhancer-promoter interactions. Analysis of single-allele multi-way interactions shows that the selected up- and downregulated gene loci are characterized by cooperative interactions between multiple enhancers and promoters, which gradually form and dissolve over the differentiation course. This indicates that enhancer-promoter hubs, which had so far only been described at high-resolution in a few selected loci^44,45^, are a common structural property of tissue-specific gene loci. Consistent with previous analyses based on multi-way 3C^45^ and imaging expriments^77^, we also observe cooperative interactions between multiple CBSs, which likely reflect stacking of CBS-anchored loops into rosette structures. Notably, we also observe multi-way interactions involving enhancers, promoters, and CBSs, which is consistent with the promiscuous interaction patterns detected by MCC. This indicates the existence of chromatin hubs involving simultaneous interactions between all classes of *cis*-regulatory elements. To better understand the role of CTCF in the formation of these hubs, we investigate the effects of CTCF depletion over the differentiation course. Interestingly, we observe that depletion of CTCF leads to a strong reduction in cooperative interactions in chromatin hubs. This shows that CTCF-dependent interactions provide a scaffold that is required for the formation of cooperative enhancer-promoter interactions in chromatin hubs (**Fig. 7**).

**Fig. 7:**
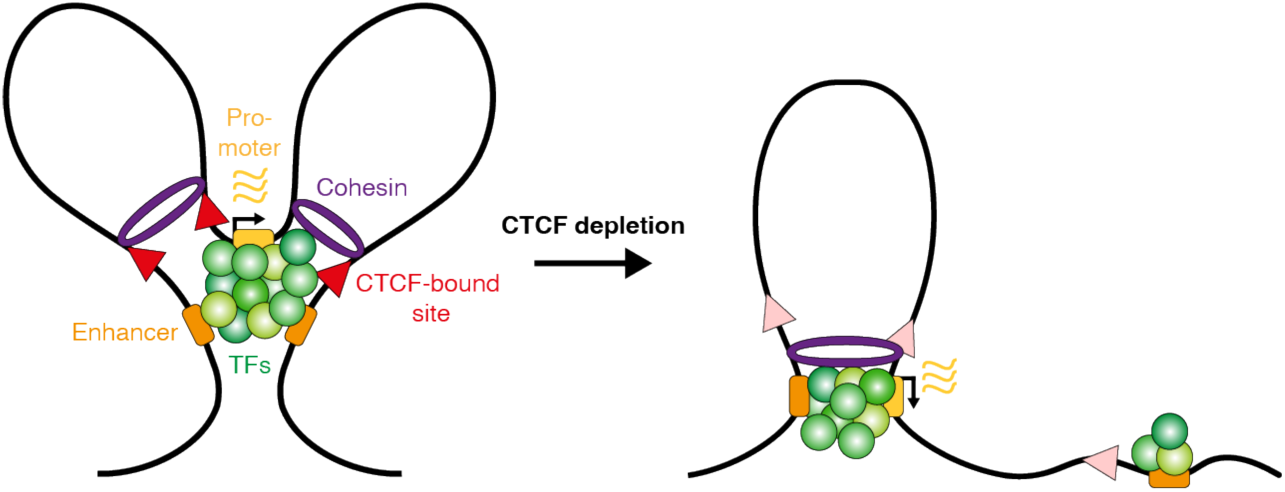
Graphical summary. Extruding cohesin molecules are stalled at CBSs, promoters, and enhancers. In presence of CTCF (left), this leads to detectable clustering of these elements in chromatin hubs. In absence of CTCF (right), these clusters form less frequently. However, enhancers still interact with their cognate promoters in a pair-wise manner (example only shown for one of the two enhancers) and thereby maintain gene expression levels. TFs = transcription factors.

In contrast to the impact of CTCF depletion on chromatin hubs, we observe that depletion of CTCF does not strongly affect pair-wise enhancer-promoter interactions. CTCF depletion therefore allows us to decouple the function of chromatin hubs and pair-wise enhancer-promoter interactions in the regulation of gene expression during cellular differentiation. Consistent with previous CTCF perturbation studies^54,73,78–83^, we do not observe widespread mis-regulation of gene expression in absence of CTCF. In most of the regions in which chromatin hubs are lost, gene expression is not significantly affected. In contrast, gene loci with significant changes in gene expression are often characterized by changes in pair-wise enhancer-promoter interactions after CTCF depletion. This indicates that pair-wise enhancer-promoter interactions have an important role in gene regulation, which is not dependent on their cooperative interactions in chromatin hubs. In agreement with our observations in the CTCF depletion experiments, we also show that changes in pair-wise enhancer-promoter interaction frequencies are strongly correlated with changes in gene expression during differentiation. Of note, the correlation coefficients that we obtain are much higher than in previous studies, which may be explained by the high resolution of our data. Since our data do not reveal any instances of pre-formed, permissive interactions, we conclude that pair-wise enhancer-promoter interactions have an instructive role in gene regulation during lymphoid-to-myeloid transdifferentiation.

Consistent with the concept of distinct classes of functional CBSs^62^, we observe that ppCBSs interact more frequently with enhancers compared to bCBSs. In addition, we observe that genes with a ppCBS are more likely to engage in long-range interactions and to be downregulated upon CTCF depletion compared to genes without a ppCBS. These and previous observations^61^ suggest that ppCBSs may contribute to the formation of (distal) enhancer-promoter interactions. However, the general importance of ppCBSs for gene regulation remains unclear since we do not observe significant changes in gene expression in many of the targeted loci that contain a ppCBS and are characterized by long-range enhancer-promoter interactions. It is possible that there are subtle changes in the expression of these genes that are difficult to detect due to previously observed increased variability of gene expression in the context of cohesin and CTCF perturbations^77^. In addition, it is of interest that we observe a tendency for proximal enhancers to interact more frequently with their cognate promoters upon CTCF depletion. Although speculative at this stage, this could provide a mechanism to compensate for loss of long-range CTCF-dependent interactions and thereby to buffer gene expression changes after CTCF depletion.

The observation that cooperative interactions in chromatin hubs are strongly reduced in absence of CTCF without a consistent effect on gene expression indicates that these hubs may not have an essential function in gene regulation. Our data therefore suggest that enhancer cooperativity is not strictly dependent on simultaneous action of enhancers at a gene promoter. This is consistent with a recent pre-print based on imaging experiments that shows that hubs are rare^84^. Although not directly tested in our study, this may have implications for the theory that transcriptional activation is dependent on the recruitment of a critical concentration of transcription factors and coactivators by clustered *cis-*regulatory elements in nuclear condensates^17–20,85^. However, it is important to note that our conclusions mostly pertain to cooperativity between enhancer clusters in a locus, since the resolution of Tri-C is not sufficient to distinguish closely spaced enhancer elements within individual enhancer clusters. It is therefore possible that higher-resolution data and/or methods that are not dependent on proximity ligation uncover cooperative interactions that are CTCF-independent and potentially more relevant for gene regulation. Furthermore, it is possible that CTCF-dependent hubs have a more pronounced function in specific cellular contexts or in regulating dynamic aspects of gene expression that are not reflected in RNA-seq data. In this regard, it is of interest that it has previously been shown that CTCF-depleted iMacs have impairments in their acute inflammatory response^54^.

Without a clear function in gene regulation, it is not obvious why CTCF-dependent chromatin hubs exist. Computational modelling of our data indicates that cooperative interactions between CBSs form as a result of cohesin-mediated loop extrusion. A relatively high density of extruding cohesin molecules and relatively stable stalling of cohesin at CBSs are in principle sufficient to explain the clustering of CBSs at the bases of multiple extruded loops, consistent with previously described rosette structures and CTCF clusters^45,77,86^. Cohesin also stalls at enhancers and promoters, which may depend on interactions with Mediator^58,71,72^ and RNA Polymerase II (RNAPII)^87–90^. This may therefore lead to clustering of enhancers and promoters with the CBS anchors. However, since cohesin stalling at enhancers and promoters is less stable compared to CBSs, cohesin-mediated loop extrusion may not lead to strong clustering of *cis*-regulatory elements in absence of the relatively stable CTCF-mediated interactions. Our model therefore suggests that chromatin hubs form as a result of the strong boundary function of CBSs during loop extrusion. Despite the minor general role of CTCF in gene regulation, it has been shown that CTCF boundaries are critical for context-specific regulation of important gene loci and that their perturbation can have severe consequences for development and disease^91–98^. It is therefore plausible that clustering of *cis*-regulatory elements in chromatin hubs arises as a by-product of strong loop extrusion boundaries, which primarily function to regulate the specificity of enhancer-promoter communication.

The development of innovative technologies to map genome architecture has led to an increasingly better understanding of the dynamic 3D structures into which the genome is organized. While some of these structures may be critical for nuclear functions, such as transcription, replication, and DNA repair, other structural features may more simply reflect the processes required to compact and organize the genome in the cell nucleus^99^. A remaining challenge therefore lies in distinguishing these two possibilities and relating genome structure to function. By combining a dynamic differentiation system, protein perturbation, and high-resolution pair-wise and multi-way analyses of chromatin interactions, our study decouples the function of pair-wise enhancer-promoter interactions from their higher-order clustering in chromatin hubs in the regulation of gene expression. Further developments in technologies to map and perturb structural and functional features of the genome will allow for more detailed dissections of their cause-consequence relationships across biological contexts in the future.

## METHODS

### Cell culture

BLaER1^52^ and BLaER1-CTCF-mAID^54^ cells were cultured in RPMI 1640 medium (Gibco, 31870025), supplemented with 10% FBS (Gibco, 10270106), 25 mM HEPES (Fisher Scientific, 15-630-080) and 2% GlutaMAX Supplement (Thermo Scientific, 35050038), at 37°C, 5% CO_2_. The cells were kept in the range of 0.2-1.5 M cells per mL and passed to fresh medium every 2 days. To transdifferentiate the B-cells into functional macrophages, fresh medium containing transdifferentiation (TD) factors (100 nM 17ß-estradiol (Calbiochem, 3301), 10 ng/mL human IL-3 (PeproTech, 200-03) and 10 ng/mL human CSF-1 (PeproTech, 300-25)) was added. The cells were differentiated up to 168 h and collected at specified time points. As a non-differentiated control (B-cell-stage, the 0 h time point), the cells were cultured in medium without TD factors. CTCF was depleted in BLaER1-CTCF-mAID cells simultaneously with inducing transdifferentiation. To achieve this, medium containing TD factors and 500 µM indole-3-acetic acid (IAA; Sigma-Aldrich, I5148), which was prepared freshly before use, were added to the cells. The cells were collected at the 96 h time-point. To ensure that IAA was not degraded, after 48 h of culture, the medium was exchanged with fresh medium supplemented with TD factors and IAA.

### Micro-Capture-C

Micro-Capture-C experiments were performed in 3 biological replicates per condition, using a procedure based on the published protocol^100^ that was further optimized for BLaER1 cells. Briefly, multiple aliquots of 15 M cells were crosslinked in 10 mL of culture medium with 2% formaldehyde (ThermoFischer, 28908). Next, the cells were permeabilized by digitonin (0.0025%; Sigma-Aldrich, D141). Digestion was performed with MNase (NEB, M0247) in low Ca^2+^ MNase buffer (10 mM Tris-HCl pH 7.5, 10 mM CaCl_2_) for 1 h at 37 °C. After confirming efficient digestion, samples were ligated and de-crosslinked. DNA was extracted using the DNeasy Blood and Tissue Kit (Qiagen, 69504). Before sonication, samples were size-selected to remove fragments corresponding to mono-nucleosomes using Mag-Bind TotalPure NGS beads (Omega Bio-Tek, M1378-01). MCC libraries were sheared using a Covaris S220 Focused-Ultrasonicator to a mean size of 200 bp (time: 280 s, duty factor: 10%, peak incident power: 175 W, cycles per burst: 200). Afterwards, the sonicated DNA fragments were indexed using 2 µg of DNA per indexing reaction. To increase library complexity, each reaction was indexed in duplicate in 7 PCR cycles using Herculase II polymerase (Agilent, 600677). The duplicate reactions were pooled and the libraires were enriched for viewpoints of interest in a double capture procedure using the KAPA Hyper Capture Reagent Kit (Roche, 9075828001). A total of 8 µg of indexed sample per biological replicate was used as input for the first capture. Oligonucleotides used for enrichment were 120 nt long and were designed with a python-based oligo tool^29^ (https://oligo.readthedocs.io/en/latest/). The oligonucleotides were ordered as a multiplexed panel of ssDNA 5′-biotinylated oligos (IDT, xGen™ Custom Hybridization Capture Panels). Each oligonucleotide of the panel was used at a final concentration of 2.9 nM. Captured DNA was pulled down using M-270 Streptavidin Dynabeads (Invitrogen, 65305), and amplified in 12 PCR cycles. All captured DNA was used as input for the second hybridization reaction. After library quality assessment with a fragment analyzer, the libraries were sequenced on the Illumina sequencing platform with 300 cycles paired-end reads.

### Capture-C and Tri-C

Capture-C and Tri-C experiments were performed in 2 and 3 biological replicates per condition, respectively, following published protocols^101,102^. Briefly, multiple aliquots of 15 M cells were crosslinked in 10 mL of culture medium, with 2% formaldehyde (ThermoFischer, 28908). The aliquots were then divided into 3 digest reactions of 5 M cells and digested with the NlaIII restriction enzyme (NEB, R0125L). After subsequent proximity ligation and DNA extraction, the 3 reactions were combined and sheared using a Covaris S220 Focused-Ultrasonicator. 6-8 µg of 3C library in 130 µL of TE buffer was sheared, to a mean size of 200 bp for Capture-C experiments (time: 180 s, duty factor: 10%, peak incident power: 175 W, cycles per burst: 200) and 450 bp for Tri-C experiments (time: 55 s, duty factor: 10%, peak incident power: 140 W, cycles per burst: 200). After sonication, Tri-C samples were size-selected with 0.7x volume of Mag-Bind TotalPure NGS beads (Omega Bio-Tek, M1378-01) for fragments longer than 300 bp. To increase library complexity, indexing was performed for each sample in duplicate (2 x 2 µg of sheared sample). Capture-C samples were indexed in 6 PCR cycles, while Tri-C samples were indexed in 7 PCR cycles. To enrich the libraries for ligated fragments containing viewpoints of interest, a double capture enrichment was performed, as described for Micro-Capture-C. The oligonucleotides used in Capture-C experiments were 70 nt long, while oligonucleotides used in Tri-C experiments were 120 nt in length. In the first enrichment round, an input of 4 µg per biological replicate was used in Capture-C experiments. To increase the complexity of enriched fragments in Tri-C experiments, an input of 9 µg per biological replicate was used. Captured DNA was pulled down using M-270 Streptavidin Dynabeads (Invitrogen, 65305), and amplified in 11 PCR cycles. All captured material was used as input for the second enrichment reaction, which was performed in the same manner. The quality of the final libraries was assessed with a fragment analyzer and the libraries were sequenced on the Illumina sequencing platform. Capture-C samples were sequenced with 150 cycles paired-end reads, while Tri-C samples were sequenced with 300 cycles paired-end reads.

### ChIPmentation

ChIPmentation experiments for CTCF were performed in 2 biological replicates per condition; ChIPmentation experiments for MED26 and SMC1A were performed in 3 biological replicates per condition during lymphoid-to-myeloid transdifferentiation and in 2 biological replicates per condition in CTCF-depleted and control cells. We followed a published protocol^55^, which we heavily optimized to improve the signal-to-noise ratio of the data. Briefly, aliquots of 1.5 M cells were crosslinked in a single fixation reaction with 1% formaldehyde (28908, Thermo Scientific) for 10 min at room temperature, quenched with ice-cold glycine (Sigma-Aldrich, G7126) at a final concentration of 125 mM, and washed 2 times with ice-cold PBS. Fixed cells were first gently lysed with Farnham lysis buffer (5 mM PIPES pH 8, 85 mM KCl, 0.5% NP-40) to isolate nuclei, which was followed by nuclear lysis with 0.5% SDS buffer (10 mM Tris-HCl pH 8, 1 mM EDTA, 0.5% SDS). Then, chromatin was fragmented, using an optimized procedure for each target. In the CTCF ChIPmentation experiments, chromatin was fragmented by sonication (time: 7 min, duty factor: 5 %, peak incident power: 140 W, cycles per burst: 200). In all MED26 and SMC1A ChIPmentation experiments, chromatin was first fragmented by a titrated amount of MNase (2.5 Kunitz) and then gently sonicated (time: 1 min, duty factor: 2 %, peak incident power: 105 W, cycles per burst: 200). To perform immunoprecipitation, a mixture of Protein A (10008D, Invitrogen) and Protein G Dynabeads (10003D, Invitrogen) in 1:1 ratio was blocked with 0.5 % BSA, washed and pre-incubated with 2 µg of primary antibody (CTCF: Diagenode, C15410210-50; SMC1A: Abcam, ab9262; MED26: Bethyl Laboratories, A302-370) and 1 µg of spike-in antibody (Biozol, 61686) for 6 h at 4 °C. Sheared chromatin was diluted in IP buffer (10 mM Tris-HCl pH 8, 1 mM EDTA, 150 mM NaCl, 1% Triton-X-100, 1X Protein Inhibitor Cocktail (Sigma-Aldrich, 11873580001)), and added along with 50 ng Drosophila spike-in chromatin to the bead-bound antibodies. The samples were incubated overnight on a rotator at 4 °C, and washed with stringent buffers the next morning. Sequencing adapters were added to the bead-bound DNA by Tagmentation with the Illumina Tagment DNA Enzyme and Buffer Kit (Illumina, 20034197). After washes, the bead-bound samples were de-crosslinked overnight at 65 °C in the presence of proteinase K (0.2 mg/mL, final) and cleaned with 1.8x volume of Mag-Bind TotalPure NGS beads (Omega Bio-Tek, M1378-01). The tagmented DNA was used for library preparation with the NEBNext High-Fidelity 2X PCR Master Mix (NEB, M0541). The quality of the libraries was assessed with a fragment analyzer and the libraries were sequenced on the Illumina sequencing platform with 75 cycles paired-end reads.

### RNA-seq

RNA-seq was performed in 2 biological replicates per condition. Briefly, cells were collected in QIAzol lysis reagent (Qiagen, 79306). RNA was isolated using phenol-chloroform extraction and the quality was assessed on a NanoDrop and fragment analyzer. 150 µg of RNA for each sample was diluted in 130 µL of RNAse-free water and sheared using a Covaris S220 Focused-Ultrasonicator (time: 10 s, duty factor: 1%, peak incident power: 100 W, cycles per burst: 200). 2 µL of sonicated RNA per sample was purified with miRNeasy Micro Kit (Qiagen, 217084) and treated with DNase I (Qiagen, 79254). 500 ng of purified RNA was used to prepare sequencing libraries with Illumina Stranded Total RNA Prep - Ligation with Ribo-Zero Plus kit (Illumina, 20040525). The quality of the libraries was assessed with a fragment analyzer and the libraries were sequenced on the Illumina sequencing platform with 100 cycles paired-end reads.

### RT-qPCR

RT-qPCR experiments were performed as previously described^53^. Briefly, cells were collected in 1 mL of QIAzol lysis reagent (Qiagen, 79306). Nucleic acids were isolated using phenol-chloroform extraction and DNA was depleted using the Turbo DNA Free Kit (ThermoFischer, AM1907) according to the manufacturer’s instructions. After conforming the quality of the RNA with an RNA ScreenTape (Agilent, 5067-5576), reverse transcription was performed with Maxima Reverse Transcriptase (ThermoScientific, EP0741) with random hexamer primers. qPCR was performed with the SYBR Select Master Mix (Applied Biosystems, 4472908) using previously validated primer sequences^52^. The data were normalized to the housekeeping gene *GAPDH*.

### Flow Cytometry

To monitor changes in cell surface markers during lymphoid-to-myeloid transdifferentiation, flow cytometry experiments were performed at different differentiation stages. Cells were harvested, washed with PBS, and blocked with human Fc Receptor Binding Antibody (eBioscience, 16-9161-73) for 20 min at 4 °C. Then, cells were stained with CD19-APC-Cy7 (BD Pharmingen, 348794) and CD11b-APC (BD Pharmingen, 550019) antibodies for 30 min at 4 °C. After washing with PBS, the cells were analyzed on a SONY SH800 cytometer. The data were analyzed with FloJo.

### Analysis of Micro-Capture-C data

Micro-Capture-C (MCC) data were processed using the MCC pipeline (hg38 genome assembly)^43^. After sequence alignment with Bowtie 2^103^, BAM files from all replicates were merged. Since capture-oligonucleotide-mediated enrichment is performed in a single hybridization reaction containing all multiplexed samples, the capture efficiency of each viewpoint is similar across samples; interactions profiles from the same viewpoint are therefore comparable across samples. To make the interaction profiles of the different viewpoints comparable, the interaction profiles were normalized for an equal number of interactions on the chromosome containing the viewpoint to correct for potential variation in the enrichment efficiency of the different capture oligonucleotides. Statistical comparisons between samples were made per viewpoint and therefore not influenced by potential variability between viewpoints. Peaks were called with MACS2^104^ (v.2.1.2), using a q-value of 0.05 and without building a shifting model. Peaks from 3 timepoints were combined and collapsed into a single unified peak set. Prior to peak annotation, the called peaks were filtered to exclude peaks outside a ± 2 Mb region surrounding the viewpoint and within ± 1 kb of the viewpoint. MCC peaks were annotated as CTCF-binding sites, promoters or enhancers based on their overlap with relevant genomic features. CTCF-binding sites were defined by overlap with CTCF ChIP-seq peaks. Promoters were defined by overlap with ATAC-seq peaks located within ± 500 bp of the transcription start site of an expressed gene. In the remaining peaks, enhancers were defined by their overlap with H3K27ac ChIP-seq peaks. Transient enhancers were defined as the enhancers of B-cell-specific genes with increased Mediator occupancy over the differentiation course. To quantify MCC interaction frequencies, MCC bigWig tracks were imported into R (v.4.2.0) and the coverage values per base-pair over MCC peaks were extracted using the R package rtracklayer (v. 1.56.1). The interaction frequency of every MCC peak was measured by calculating the average of the highest 70% of base-pair bigWig coverage values. For quantifications of the proportion of interactions of promoter, ppCBS, and bCBS viewpoints with *cis-*regulatory elements across the targeted loci (**Fig. 2f**), CBS viewpoints were classified as ppCBS (and not bCBS) in case they were proximal to both a promoter and a boundary. Similar to quantification of the MCC signal, the occupancy of SMC1A, MED26, and CTCF at annotated MCC peaks was quantified by extracting the coverage information from the corresponding bigWig files and calculating the average of the highest 70% of base-pair coverage values across the MCC peaks.

### Analysis of Capture-C and Tri-C data

Capture-C and Tri-C data were analyzed using the CapCruncher pipeline^101^ (v.0.2.3) in capture mode (hg38 genome assembly). Data from all replicates were merged. Capture-C data were normalized as described for MCC. For Tri-C analysis, a custom script was used to extract reads with 2 or more *cis* reporters. These reads were used to calculate multi-way interaction counts between reporter fragments for each viewpoint. Contact matrices in which multiple 3-way interactions were calculated from reads containing more than 3 reporters did not show any qualitative differences compared to contact matrices in which only a single 3-way interaction per read was included. All detected 3-way interactions were therefore included in the contact matrices to boost the complexity of the data. The interaction counts were binned in 1500-4000 bp bins and corrected for the number of restriction fragments present in each bin. The matrices were further normalized for the total multi-way interaction counts within a 2 Mb region (which was corrected for the number of restriction fragments present in the region) surrounding the viewpoint. To quantify the interaction frequencies between two regions of interest (ROIs), a 3X3 binning matrix surrounding the pair of ROIs was extracted from the normalized contact matrices and averaged.

### Analysis of ChIPmentation data

ChIPmentation data were analyzed with the NGseqBasic pipeline^12^ (hg38 genome assembly). After sequence alignment and deduplication, bigWig coverage tracks were generated with deepTools (v.3.3.0) bamCoverage function, using a binning size of 10 bp. CTCF ChIPmentation coverage tracks were normalized using spike-in reads count, while SMC1A and MED26 coverage tracks were normalized using RPKM. Coverage tracks of replicates were merged using UCSC bigWigMerge (v.2).

### Analysis of RNA-seq data

RNA-seq data were processed using the nf-core/rnaseq pipeline (v.3.12.0; hg38 genome assembly). After sequence alignment with STAR^105^ (v.2.6.1d), the expression of protein-coding exons was quantified with featureCounts (v.2.0.6). DESeq2^106^ (v.1.36.0) was used to perform differential gene expression analysis, with a threshold of adjusted p-value < 0.01 and absolute log2 fold change (FC) > 0.6. To compare the genomic landscapes and enhancer distributions of differentially expressed genes with those of stable genes upon CTCF depletion, promoters and enhancers were paired based on the previously described TAD pairing strategy^53^, in which promoters and enhancers within the same TAD are paired if the Spearman’s correlation coefficient between mRNA and eRNA read counts during transdifferentiation is higher than 0.4. The distances between all paired promoters and enhancers were plotted in the distance distributions (**Extended Data Figure 6d**).

### Correlation analyses

Spearman rank correlation was used to measure the association between changes in pair-wise chromatin interactions and changes in various features of interest, including SMC1A, MED26, and CTCF occupancy, H3K27ac levels, and eRNA expression. Changes in interaction frequencies and different signals at corresponding MCC peaks were calculated at 24h and 96h relative to the baseline at 0h. To correlate pair-wise enhancer-promoter interaction frequencies and gene expression levels, a total enhancer-promoter interaction score was defined for every targeted promoter as the sum of the signal of the enhancer peaks in the MCC interaction profiles. For all targeted loci, changes in gene expression levels, total enhancer-promoter interaction scores, and the products of the total enhancer-promoter interaction scores and promoter accessibility, based on TT-seq, MCC, and ATAC-seq data, were calculated. Changes at 24 h and 96 h were calculated relative to the baseline at 0 h. Spearman rank correlation coefficients were computed to investigate the relationship between changes in pair-wise enhancer-promoter interactions and changes in gene expression levels during lymphoid-to-myeloid differentiation, as well as the relationship between changes in MED26 and SMC1A occupancy and pair-wise enhancer-promoter interactions after CTCF depletion.

### Molecular dynamics simulations

2 Mb regions around the *CCR1*, *NFKBIZ*, and *TRIB1* loci were modelled based on a previously described modelling framework^66,67^ using the multipurpose EspressoMD package^107^. Briefly, the chromatin fiber is modelled as a self-avoiding polymer chain consisting of equisized beads representing 2 kb of chromatin. Beads were classified as binding the transcription machinery (i.e., promoters and enhancers; based on Mediator ChIPmentation peaks), binding CTCF (based on CTCF ChIPmentation peaks, including motif orientation), as transcribed genic regions (gene bodies of active genes; based on TT-seq data), or as neutral (with none of the above-mentioned features). Polymers were used in molecular dynamics simulations in a 3D space following Langevin equations to model thermal motion of chromatin and its binding factors in an implicit solvent (the nucleoplasm), with the following postulations^66^: (1) the transcription machinery has affinity for promoters and enhancers and traverses gene bodies; (2) components of the transcription machinery (RNAPII, Mediator) also have affinity for each other to simulate condensate formation; (3) cohesin complexes can bind anywhere on the polymer and extrude loops, but have a preference for loading at enhancers and promoters. To account for the experimentally observed accumulation of cohesin at CTCF- and Mediator-bound sites, extrusion of loops by cohesin stalls when encountering a CTCF-bound site with a motif oriented towards the direction of extrusion or when meeting the transcription machinery. By dynamically forming and dissolving protein-chromatin bonds, this framework simulates chromatin loop extrusion by the cohesin complex and traversing of RNAPII during transcription. The rates of loop extrusion and RNAPII translocation along chromatin were set within the range of experimentally deduced values (RNAPII: 1-5 kb/min; cohesin: 15-30 kb/min)^67^. RNAPII-cohesin and cohesin-cohesin crossing rates were set at 1.5 and 0.15 crossings per second, respectively.

The ensemble of chromatin conformations resulting from these simulations were used to generate pair-wise interaction profiles and three-way contact matrices among the beads of each polymer, allowing for direct quantitative comparisons between the model and the experimental MCC and Tri-C data, respectively. The degree of triplet colocalization of the viewpoint with regions A and B, over what would be expected by the probability of independent, pair-wise interactions (**Extended Data Fig. 5**), was estimated from the predicted structures using the correlation coefficient:

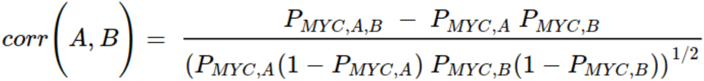

### Analysis of reference datasets

Available ATAC-seq^54^ (GSE131620), CTCF ChIP-seq^54^ (GSE140528), and H3K27ac ChIP-seq^54^ (E-MTAB-9010) data were analyzed with the NGseqBasic pipeline^108^ (hg38 genome assembly). ATAC-seq and H3K27ac ChIP-seq peaks were identified using MACS2^104^ (v.2.1.2) in broad peak calling mode with a q-value of 0.01 and a broad cutoff of 0.05. Peaks in CTCF ChIP-seq data were identified using MACS2 in narrow peak calling mode with a q-value of 0.01. H3K27me3 ChIP-seq bigWig tracks (GSE259000, GSE256663 and GSE257306) were downloaded from ENCODE. Hi-C data^54^ (GSE131620) were analyzed with Hi-C Pro^109^ (Servant, 2015) and plotted using cooltools^110^. TT-seq data^53^ (GSE131620) were processed with the nf-core/rnaseq pipeline as described for RNA-seq data analysis. To quantify mRNA expression levels, featureCounts was used to count reads over the exons of all protein-coding genes. To quantify eRNA expression levels, featureCounts was used to count reads over the MCC enhancer peak regions. The raw count data were normalized by DESeq2-estimated size factors.

## Data availability

The Micro-Capture-C, Capture-C, Tri-C, ChIPmentation and RNA-seq datasets generated and analyzed for the current study are available from the Gene Expression Omnibus (GEO) as a SuperSeries under accession number GSE263641.

## ACKNOWLEDGEMENTS

We would like to thank T. Graf (Centre for Genomic Regulation, Barcelona, Spain) and G. Stik (Josep Carreras Leukaemia Research Institute, Barcelona, Spain) for providing the BLaER1 and BLaER1-CTCF-mAID cell lines. We are grateful to P. Cramer for advice, discussions, and infrastructure support. We are also grateful for support from the Facility for Light Microscopy at the Max Planck Institute for Multidisciplinary Sciences, particularly from Jasmin Jakobi, Peter Lénart, and Antonio Politi. We would like to thank J. Söding for advice on bioinformatic analysis and J. Choi, K. Lysakovskaia, K. Maier, M. Rohm, P. Rus, G. Stik and K. Zumer for experimental advice and support. We are also grateful to K. Schmitt for support with data visualization and members of the Oudelaar group for helpful discussions and feedback. This work was supported by the Max Planck Society (A.M.O); the Deutsche Forschungsgemeinschaft (DFG) via SFB 1565 (project 469281184 / P02 to A.M.O. and project 469281184 / P03 to A.P.), SPP 2202 (project 507778679 to A.M.O. and project 422389065 to A.P.) and SPP 2191 (project 506296585 to A.P.); the PhD program "Genome Science" – International Max Planck Research School at the Georg August University Göttingen (M.A.K., S.R., T.B.N.C., A.A.); and the MSc/PhD program "Molecular Biology" – International Max Planck Research School at the Georg August University Göttingen (Y.Z., Z.F.G., N.V.).

## AUTHOR CONTRIBUTIONS

M.A.K. carried out most of the experiments, performed basic data analyses, and prepared the figures. Y.Z. performed the majority of the bioinformatic analyses. Z.F.G. performed MCC experiments over the differentiation course. S.R. optimized the ChIPmentation protocol and performed ChIPmentation experiments. M.B. performed molecular dynamics simulations. T.B.N.C. performed RNA-seq experiments. N.V. performed ChIPmentation experiments and bioinformatic analyses. A.A. performed ChIPmentation experiments. M.L. assisted with bioinformatic analyses. A.P. provided conceptual advice, supervised the molecular dynamics simulations, and acquired funding. A.M.O. conceived and supervised the project, acquired funding, and wrote the manuscript, with input from M.A.K., Y.Z., M.B., and A.P..

## REFERENCES

1. Long, H.K., Prescott, S.L. & Wysocka, J. Ever-Changing Landscapes: Transcriptional Enhancers in Development and Evolution. Cell 167, 1170–1187 (2016).

2. Cramer, P. Organization and regulation of gene transcription. Nature 573, 45–54 (2019).

3. Oudelaar, A.M. & Higgs, D.R. The relationship between genome structure and function. Nature Reviews Genetics 22, 154–168 (2021).

4. Schoenfelder, S. & Fraser, P. Long-range enhancer–promoter contacts in gene expression control. Nature Reviews Genetics 20, 437–455 (2019).

5. Furlong, E.E.M. & Levine, M. Developmental enhancers and chromosome topology. Science 361, 1341–1345 (2018).

6. Kim, J. & Dean, A. Enhancers navigate the three-dimensional genome to direct cell fate decisions. Current Opinion in Structural Biology 71, 101–109 (2021).

7. Ibrahim, D.M. & Mundlos, S. The role of 3D chromatin domains in gene regulation: a multi-facetted view on genome organization. Current Opinion in Genetics & Development 61, 1–8 (2020).

8. Symmons, O. et al. Functional and topological characteristics of mammalian regulatory domains. Genome Research 24, 390–400 (2014).

9. Nora, E.P. et al. Spatial partitioning of the regulatory landscape of the X-inactivation centre. Nature 485, 381–385 (2012).

10. Dixon, J.R. et al. Topological domains in mammalian genomes identified by analysis of chromatin interactions. Nature 485, 376–380 (2012).

11. Fudenberg, G. et al. Formation of Chromosomal Domains by Loop Extrusion. CellReports 15, 2038–2049 (2016).

12. Sanborn, A.L. et al. Chromatin extrusion explains key features of loop and domain formation in wild-type and engineered genomes. Proceedings of the National Academy of Sciences of the United States of America 112, E6456–65 (2015).

13. Karpinska, M.A. & Oudelaar, A.M. The role of loop extrusion in enhancer-mediated gene activation. Current Opinion in Genetics & Development 79, 102022 (2023).

14. de Wit, E. & Nora, E.P. New insights into genome folding by loop extrusion from inducible degron technologies. Nature Reviews Genetics (2022).

15. Stadhouders, R., Filion, G.J. & Graf, T. Transcription factors and 3D genome conformation in cell-fate decisions. Nature 569, 345–354 (2019).

16. Kim, S. & Shendure, J. Mechanisms of Interplay between Transcription Factors and the 3D Genome. Molecular Cell 76, 306–319 (2019).

17. Hnisz, D., Shrinivas, K., Young, R.A., Chakraborty, A.K. & Sharp, P.A. A Phase Separation Model for Transcriptional Control. Cell 169, 13–23 (2017).

18. Boija, A. et al. Transcription Factors Activate Genes through the Phase-Separation Capacity of Their Activation Domains. Cell 175, 1842–1855.e16 (2018).

19. Sabari, B.R. et al. Coactivator condensation at super-enhancers links phase separation and gene control. Science 361, eaar3958 (2018).

20. Cho, W.-K. et al. Mediator and RNA polymerase II clusters associate in transcription-dependent condensates. Science 361, 412 (2018).

21. Mir, M., Bickmore, W., Furlong, E.E.M. & Narlikar, G. Chromatin topology, condensates and gene regulation: shifting paradigms or just a phase? Development 146, dev182766 (2019).

22. Ghavi-Helm, Y. et al. Enhancer loops appear stable during development and are associated with paused polymerase. Nature 513, 89–100 (2014).

23. Chen, Z. et al. Increased enhancer–promoter interactions during developmental enhancer activation in mammals. Nature Genetics 56, 675–685 (2024).

24. Espinola, S.M. et al. Cis-regulatory chromatin loops arise before TADs and gene activation, and are independent of cell fate during early Drosophila development. Nature Genetics 53, 477–486 (2021).

25. Hug, C.B., Grimaldi, A.G., Kruse, K. & Vaquerizas, J.M. Chromatin Architecture Emerges during Zygotic Genome Activation Independent of Transcription. Cell 169, 216-228.e19 (2017).

26. Pollex, T. et al. Enhancer–promoter interactions become more instructive in the transition from cell-fate specification to tissue differentiation. Nature Genetics (2024).

27. Murphy, D. et al. 3D Enhancer–promoter networks provide predictive features for gene expression and coregulation in early embryonic lineages. Nature Structural & Molecular Biology 31, 125–140 (2024).

28. Andrey, G. et al. Characterization of hundreds of regulatory landscapes in developing limbs reveals two regimes of chromatin folding. Genome Research 27, 223–233 (2017).

29. Oudelaar, A.M. et al. Dynamics of the 4D genome during in vivo lineage specification and differentiation. Nature Communications 11, 2722 (2020).

30. Beagan, J.A. et al. Three-dimensional genome restructuring across timescales of activity-induced neuronal gene expression. Nature Neuroscience 23, 707–717 (2020).

31. Bonev, B. et al. Multiscale 3D Genome Rewiring during Mouse Neural Development. Cell 171, 557–572.e24 (2017).

32. Caputo, L. et al. The Isl1/Ldb1 Complex Orchestrates Genome-wide Chromatin Organization to Instruct Differentiation of Multipotent Cardiac Progenitors. Cell Stem Cell 17, 287–299 (2015).

33. Phanstiel, D.H. et al. Static and Dynamic DNA Loops form AP-1-Bound Activation Hubs during Macrophage Development. Molecular Cell 67, 1037–1048.e6 (2017).

34. Siersbæk, R. et al. Dynamic Rewiring of Promoter-Anchored Chromatin Loops during Adipocyte Differentiation. Molecular Cell 66, 420–435.e5 (2017).

35. Rubin, A.J. et al. Lineage-specific dynamic and pre-established enhancer–promoter contacts cooperate in terminal differentiation. Nature Genetics 49, 1522–1528 (2017).

36. Dall’Agnese, A. et al. Transcription Factor-Directed Re-wiring of Chromatin Architecture for Somatic Cell Nuclear Reprogramming toward trans-Differentiation. Molecular Cell 76, 453–472.e8 (2019).

37. Stadhouders, R. et al. Transcription factors orchestrate dynamic interplay between genome topology and gene regulation during cell reprogramming. Nature Genetics 50, 238–249 (2018).

38. de Laat, W. & Duboule, D. Topology of mammalian developmental enhancers and their regulatory landscapes. Nature 502, 499–506 (2013).

39. Xiao, J.Y., Hafner, A. & Boettiger, A.N. How subtle changes in 3D structure can create large changes in transcription. eLife 10, e64320 (2021).

40. Zuin, J. et al. Nonlinear control of transcription through enhancer–promoter interactions. Nature 604, 571–577 (2022).

41. Dekker, J., Rippe, K., Dekker, M. & Kleckner, N. Capturing chromosome conformation. Science 295, 1306–1311 (2002).

42. Davies, J.O.J., Oudelaar, A.M., Higgs, D.R. & Hughes, J.R. How best to identify chromosomal interactions: a comparison of approaches. Nature Methods 14, 125–134 (2017).

43. Hua, P. et al. Defining genome architecture at base-pair resolution. Nature 595, 125–129 (2021).

44. Oudelaar, A.M. et al. Single-allele chromatin interactions identify regulatory hubs in dynamic compartmentalized domains. Nature Genetics 50, 1744–1751 (2018).

45. Allahyar, A. et al. Enhancer hubs and loop collisions identified from single-allele topologies. Nature Genetics 50, 1151–1160 (2018).

46. Chang, L.-H. et al. Multi-feature clustering of CTCF binding creates robustness for loop extrusion blocking and Topologically Associating Domain boundaries. Nature Communications 14, 5615 (2023).

47. Deshpande, A.S. et al. Identifying synergistic high-order 3D chromatin conformations from genome-scale nanopore concatemer sequencing. Nature Biotechnology (2022).

48. Zhong, J.-Y. et al. High-throughput Pore-C reveals the single-allele topology and cell type-specificity of 3D genome folding. Nature Communications 14, 1250 (2023).

49. Olivares-Chauvet, P. et al. Capturing pairwise and multi-way chromosomal conformations using chromosomal walks. Nature 540, 296–300 (2016).

50. Quinodoz, S.A. et al. Higher-Order Inter-chromosomal Hubs Shape 3D Genome Organization in the Nucleus. Cell 174, 744–757.e24 (2018).

51. Beagrie, R.A. et al. Complex multi-enhancer contacts captured by genome architecture mapping. Nature 543, 519–524 (2017).

52. Rapino, F. et al. C/EBPA Induces Highly Efficient Macrophage Transdifferentiation of B Lymphoma and Leukemia Cell Lines and Impairs Their Tumorigenicity. Cell Reports 3, 1153–1163 (2013).

53. Choi, J. et al. Evidence for additive and synergistic action of mammalian enhancers during cell fate determination. eLife 10, e65381 (2021).

54. Stik, G. et al. CTCF is dispensable for immune cell transdifferentiation but facilitates an acute inflammatory response. Nature Genetics 52, 655–661 (2020).

55. Schmidl, C., Rendeiro, A.F., Sheffield, N.C. & Bock, C. ChIPmentation: fast, robust, low-input ChIP-seq for histones and transcription factors. Nature Methods 12, 963–965 (2015).

56. Satoh, T. et al. Critical role of Trib1 in differentiation of tissue-resident M2-like macrophages. Nature 495, 524–528 (2013).

57. Wendt, K.S. et al. Cohesin mediates transcriptional insulation by CCCTC-binding factor. Nature 451, 796–801 (2008).

58. Kagey, M.H. et al. Mediator and cohesin connect gene expression and chromatin architecture. Nature 467, 430–435 (2010).

59. Thomas, M.D., Kremer, C.S., Ravichandran, K.S., Rajewsky, K. & Bender, T.P. c-Myb Is Critical for B Cell Development and Maintenance of Follicular B Cells. Immunity 23, 275–286 (2005).

60. Vermunt, M.W. et al. Gene silencing dynamics are modulated by transiently active regulatory elements. Molecular Cell 83, 715–730.e6 (2023).

61. Kubo, N. et al. Promoter-proximal CTCF binding promotes distal enhancer-dependent gene activation. Nature Structural & Molecular Biology 28, 152–161 (2021).

62. Huang, H. et al. CTCF mediates dosage- and sequence-context-dependent transcriptional insulation by forming local chromatin domains. Nature Genetics 53, 1064–1074 (2021).

63. Oeckinghaus, A., Hayden, M.S. & Ghosh, S. Crosstalk in NF-κB signaling pathways. Nature Immunology 12, 695–708 (2011).

64. Kaufmann, A., Salentin, R., Gemsa, D. & Sprenger, H. Increase of CCR1 and CCR5 expression and enhanced functional response to MIP-1α during differentiation of human monocytes to macrophages. Journal of Leukocyte Biology 69, 248–252 (2001).

65. Kuo, T.C. & Schlissel, M.S. Mechanisms controlling expression of the RAG locus during lymphocyte development. Current Opinion in Immunology 21, 173–178 (2009).

66. Zhang, S., Übelmesser, N., Barbieri, M. & Papantonis, A. Enhancer–promoter contact formation requires RNAPII and antagonizes loop extrusion. Nature Genetics (2023).

67. Platania, A. et al. Transcription processes compete with loop extrusion to homogenize promoter and enhancer dynamics. Science Advances 10, eadq0987 (2024).

68. Nikolic, T. et al. The DNA-binding factor Ctcf critically controls gene expression in macrophages. Cellular & Molecular Immunology 11, 58–70 (2014).

69. Luan, J., et al. Distinct properties and functions of CTCF revealed by a rapidly inducible degron system. Cell Reports 34(2021).

70. Merkenschlager, M. & Nora, E .P. CTCF and Cohesin in Genome Folding and Transcriptional Gene Regulation. dx.doi.org 17, 17–43 (2016).

71. Phillips-Cremins, J.E. et al. Architectural Protein Subclasses Shape 3D Organization of Genomes during Lineage Commitment. Cell 153, 1281–1295 (2013).

72. Ramasamy, S. et al. The Mediator complex regulates enhancer-promoter interactions. Nature Structural & Molecular Biology 30, 991–1000 (2023).

73. Aljahani, A. et al. Analysis of sub-kilobase chromatin topology reveals nano-scale regulatory interactions with variable dependence on cohesin and CTCF. Nature Communications 13, 2139 (2022).

74. Kane, L. et al. Cohesin is required for long-range enhancer action at the Shh locus. Nature Structural & Molecular Biology 29, 891–897 (2022).

75. Rinzema, N.J. et al. Building regulatory landscapes reveals that an enhancer can recruit cohesin to create contact domains, engage CTCF sites and activate distant genes. Nature Structural & Molecular Biology 29, 563–574 (2022).

76. Calderon, L. et al. Cohesin-dependence of neuronal gene expression relates to chromatin loop length. eLife 11, e76539 (2022).

77. Hafner, A. et al. Loop stacking organizes genome folding from TADs to chromosomes. Molecular Cell 83, 1377–1392.e6 (2023).

78. Nora, E.P. et al. Targeted Degradation of CTCF Decouples Local Insulation of Chromosome Domains from Genomic Compartmentalization. Cell 169, 930–944.e22 (2017).

79. Wutz, G. et al. Topologically associating domains and chromatin loops depend on cohesin and are regulated by CTCF, WAPL, and PDS5 proteins. The EMBO Journal 36, 3573–3599 (2017).

80. Despang, A. et al. Functional dissection of the Sox9–Kcnj2 locus identifies nonessential and instructive roles of TAD architecture. Nature Genetics 51, 1263–1271 (2019).

81. Hsieh, T.-H.S. et al. Enhancer–promoter interactions and transcription are largely maintained upon acute loss of CTCF, cohesin, WAPL or YY1. Nature Genetics 54, 1919–1932 (2022).

82. Goel, V.Y., Huseyin, M.K. & Hansen, A.S. Region Capture Micro-C reveals coalescence of enhancers and promoters into nested microcompartments. bioRxiv, 2022.07.12.499637 (2022).

83. Liu, Y. et al. CTCF coordinates cell fate specification via orchestrating regulatory hubs with pioneer transcription factors. Cell Reports 42, 113259 (2023).

84. Le, D.J., Hafner, A., Gaddam, S., Wang, K.C. & Boettiger, A.N. Super-enhancer interactomes from single cells link clustering and transcription. bioRxiv, 2024.05.08.593251 (2024).

85. Lee, R. et al. CTCF-mediated chromatin looping provides a topological framework for the formation of phase-separated transcriptional condensates. Nucleic Acids Research 50, 207–226 (2022).

86. Gu, B. et al. Opposing Effects of Cohesin and Transcription on CTCF Organization Revealed by Super-resolution Imaging. Molecular Cell 80, 699–711.e7 (2020).

87. Zhang, S., Übelmesser, N., Barbieri, M. & Papantonis, A. Enhancer–promoter contact formation requires RNAPII and antagonizes loop extrusion. Nature Genetics 55, 832–840 (2023).

88. Barshad, G. et al. RNA polymerase II dynamics shape enhancer–promoter interactions. Nature Genetics 55, 1370–1380 (2023).

89. Valton, A.-L. et al. A cohesin traffic pattern genetically linked to gene regulation. bioRxiv, 2021.07.29.454218 (2021).

90. Busslinger, G.A. et al. Cohesin is positioned in mammalian genomes by transcription, CTCF and Wapl. Nature 544, 503–507 (2017).

91. Franke, M. et al. Formation of new chromatin domains determines pathogenicity of genomic duplications. Nature 538, 265–269 (2016).

92. Lupiáñez, D.G. et al. Disruptions of Topological Chromatin Domains Cause Pathogenic Rewiring of Gene-Enhancer Interactions. Cell 161, 1012–1025 (2015).

93. Narendra, V. et al. CTCF establishes discrete functional chromatin domains at the Hox clusters during differentiation. Science 347, 1017–1021 (2015).

94. Flavahan, W.A. et al. Insulator dysfunction and oncogene activation in IDH mutant gliomas. Nature 529, 110–114 (2016).

95. Hnisz, D. et al. Activation of proto-oncogenes by disruption of chromosome neighborhoods. Science 351, 1454–1458 (2016).

96. Dowen, J.M. et al. Control of Cell Identity Genes Occurs in Insulated Neighborhoods in Mammalian Chromosomes. Cell 159, 374–387 (2014).

97. Hanssen, L.L.P. et al. Tissue-specific CTCF–cohesin-mediated chromatin architecture delimits enhancer interactions and function in vivo. Nature Cell Biology 19, 952–961 (2017).

98. Anania, C. et al. In vivo dissection of a clustered-CTCF domain boundary reveals developmental principles of regulatory insulation. Nature Genetics 54, 1026–1036 (2022).

99. Solovei, I. & Mirny, L. Spandrels of the cell nucleus. Current Opinion in Cell Biology 90, 102421 (2024).

100. Hamley, J.C., Li, H., Denny, N., Downes, D. & Davies, J.O.J. Determining chromatin architecture with Micro Capture-C. Nature Protocols 18, 1687–1711 (2023).

101. Downes, D.J. et al. Capture-C: a modular and flexible approach for high-resolution chromosome conformation capture. Nature Protocols 17, 445–475 (2022).

102. Oudelaar, A.M., Downes, D.J. & Hughes, J.R. Assessment of Multiway Interactions with Tri-C. in Spatial Genome Organization: Methods and Protocols (ed. Sexton, T.) 95–112 (Springer US, New York, NY, 2022).

103. Langmead, B. & Salzberg, S.L. Fast gapped-read alignment with Bowtie 2. Nature Methods 9, 357–359 (2012).

104. Zhang, Y. et al. Model-based analysis of ChIP-Seq (MACS). Genome biology 9, R137 (2008).

105. Dobin, A. et al. STAR: ultrafast universal RNA-seq aligner. Bioinformatics 29, 15–21 (2013).

106. Love, M.I., Huber, W. & Anders, S. Moderated estimation of fold change and dispersion for RNA-seq data with DESeq2. Genome biology 15, 550 (2014).

107. Reynwar, B.J. et al. Aggregation and vesiculation of membrane proteins by curvature-mediated interactions. Nature 447, 461–464 (2007).

108. Telenius, J., The, W.C. & Hughes, J.R. NGseqBasic - a single-command UNIX tool for ATAC-seq, DNaseI-seq, Cut-and-Run, and ChIP-seq data mapping, high-resolution visualisation, and quality control. bioRxiv, 393413 (2018).

109. Servant, N. et al. HiC-Pro: an optimized and flexible pipeline for Hi-C data processing. Genome biology 16, 259 (2015).

110. Open2C et al. Cooltools: enabling high-resolution Hi-C analysis in Python. bioRxiv, 2022.10.31.514564 (2022).

