## Supplementary Figures for "CTCF depletion decouples enhancer-mediated gene activation from chromatin hub formation during cellular differentiation"

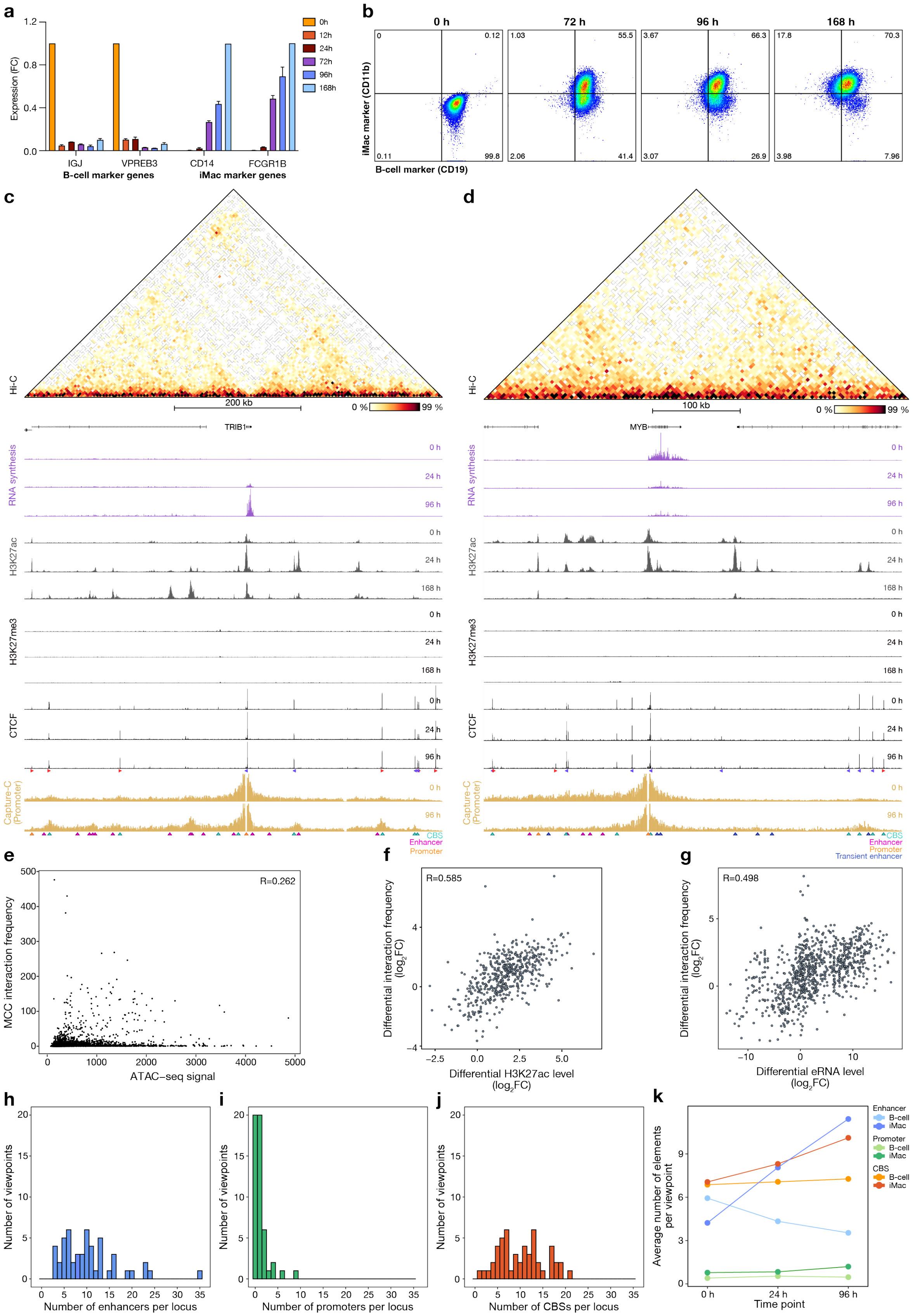


***Extended Data Fig. 1: Characterization of the BLaER1 lymphoid-to-myeloid transdifferentiation system.*** *(a) Changes in the expression levels of B-cell and iMac marker genes during lymphoid-to-myeloid transdifferentiation, measured with RT-qPCR at 0 h, 12 h, 24 h, 72 h, 96 h, and 168 h after differentiation induction. The expression of B-cell marker genes (IGJ, VPREB3) is shown as fold change (FC) relative to the 0 h timepoint, whereas the expression of iMac marker genes (CD14, FCGR1B) is shown relative to 168 h. The bars represent the averages*

*of n = 2 replicates; the error bars indicate the standard deviation. (b) Changes in cell surface markers during lymphoid-to-myeloid transdifferentiation, measured by flow cytometry. A representative example of n = 2 biological replicates is shown. (c) Chromatin landscape of the TRIB1 locus (chr8:125,079,965-125,739,965; 660 kb) during lymphoid-to-myeloid transdifferentiation. From top to bottom: Hi-C contact matrix (5 kb resolution) at 96 h; gene annotation; RNA synthesis (TT-seq) at 0 h, 24 h, and 96 h; H3K27ac occupancy (H3K27ac ChIP-seq) at 0 h, 24 h, and 168 h; H3K27me3 occupancy (H3K27me3 ChIP-seq) at 0 h, 24 h, and 168 h; CTCF occupancy (CTCF ChIP-seq) at 0 h, 24 h, and 96 h; Capture-C data from the viewpoint of the TRIB1 promoter at 0 h and 96 h. The axes of the profiles are scaled to signal and have the following ranges: RNA synthesis = 0–10455; H3K27ac = 0–5311; CTCF = 0–12178; Capture-C = 0–1800. The orientations of CTCF motifs at prominent CTCF-binding sites are indicated by arrowheads (forward orientation in red; reverse orientation in blue). Interactions detected by MCC (Fig. 1) with CBSs, enhancers, and promoters are annotated with cyan, magenta, and orange triangles, respectively. (d) Chromatin landscape of the MYB locus (chr6:134,992,474-135,472,474; 480 kb) during lymphoid-to-myeloid transdifferentiation, as described in panel c, except that Hi-C data are shown at 0 h and that MCC interactions with transient enhancers are annotated with blue triangles. The axes of the profiles are scaled to signal and have the following ranges: RNA synthesis = 0–12791; H3K27ac = 0–3930; CTCF = 0–6052; Capture-C = 0–2500. (e) Correlation between ATAC-seq signals (± 500 kb from the viewpoint) and MCC interaction frequencies, based on Spearman's correlation test. (f) Correlation between differential enhancer-promoter interaction frequencies and differential H3K27ac levels at the interacting elements (24 h vs 0h), based on Spearman's correlation test. (g) Correlation between differential enhancer-promoter interaction frequencies and differential eRNA levels at the interacting elements (24 h vs 0h and 96 h vs 0 h), derived from TT-seq, as described in panel f. (h) Histogram showing the distribution of the number of enhancer interactions with the promoter viewpoints per locus as detected by MCC. (i) Histogram showing the distribution of the number of promoter interactions with the promoter viewpoints per locus as detected by MCC. (j) Histogram showing the distribution of the number of CBS interactions with the promoter viewpoints per locus as detected by MCC. (k) Overview of the average number of enhancer, promoter, and CBS interactions with the promoter viewpoints as detected by MCC at 0 h, 24 h, and 96 h.*

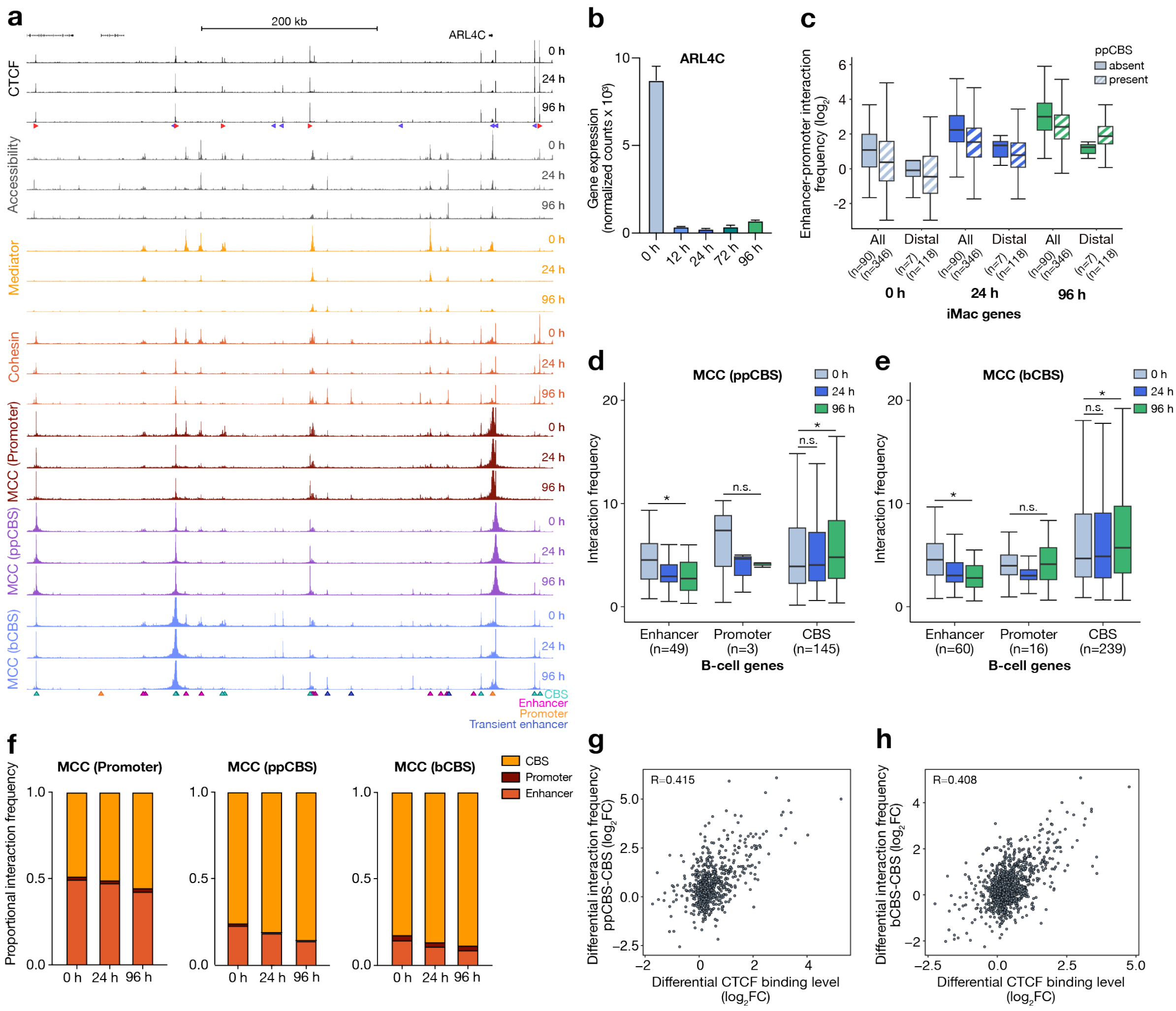


***Extended Data Fig. 2: Interaction patterns of promoters and CTCF-binding sites through lymphoid-to-myeloid transdifferentiation.*** *(a) Chromatin landscape of the ARLC4 locus (chr2:233,966,201-234,566,201; 600 kb) during lymphoid-to-myeloid transdifferentiation. From top to bottom: gene annotation; CTCF occupancy (CTCF ChIP-seq) at 0 h, 24 h, and 96 h; chromatin accessibility (ATAC-seq) at 0 h, 24 h, and 96 h; Mediator occupancy (MED26 ChIPmentation) at 0 h, 24 h, and 96 h; Cohesin occupancy (SMC1A ChIPmentation) at 0 h, 24 h, and 96 h; Micro-Capture-C (MCC) data from the viewpoint of the promoter, promoter-proximal CTCF-binding site (ppCBS) and boundary CTCF-binding site (bCBS) at 0 h, 24 h, and 96 h. The axes of the profiles are scaled to signal and have the following ranges: CTCF = 0–9200; Accessibility = 0–3204; Mediator = 0–5636; Cohesin = 0–2515. The orientations of CTCF motifs at prominent CBSs are indicated by arrowheads (forward orientation in red; reverse orientation in blue). MCC interactions with CBSs, enhancers, promoters, and transient enhancers are annotated with cyan, magenta, orange, and blue triangles, respectively. (b) ARLC4 expression at 0 h, 12 h, 24 h, 72 h, and 96 h after differentiation induction. Expression levels are derived from TT-seq data. The bars represent the average of n = 2 replicates; the error bars indicate the standard deviation. (c) Comparison of enhancer-promoter interaction frequencies of iMac-specific genes with and without a ppCBS (± 5 kb from the promoter) at 0 h, 24 h, and 96 h after differentiation induction. The analysis is performed for all interacting enhancers and distal (> 150 kb) enhancers only and shows that iMac genes with a ppCBS have lower baseline interactions with all enhancers compared to genes without a ppCBS but increased interactions specifically with distal enhancers at 96 h. (d) Interaction frequencies of ppCBSs of B-cell-specific genes with enhancers, promoters, and CBSs at 0 h, 24 h, and 96 h after differentiation induction. Boxplots show the interquartile range (IQR) and median of the data; whiskers indicate the minima and maxima within 1.5 * IQR; asterisks indicate significance ($P < 0.01$, two-sided paired Wilcoxon signed rank test). (e) Interaction frequencies of bCBSs of B-cell-specific genes, as described in panel d. (f) Comparison of the proportion of interactions of promoters, ppCBSs and bCBSs with enhancers, promoters, and CBSs in B-cell-specific gene loci at 0 h, 24 h, and 96 h. (g) Correlation between differential interaction frequencies between ppCBS and CBSs and differential CTCF binding levels at the interacting elements (24 h vs 0h and 96 h vs 0 h), based on Spearman's correlation test. (h) Correlation between differential interaction frequencies between bCBS and CBSs and differential CTCF binding levels at the interacting elements, as described in panel g.*

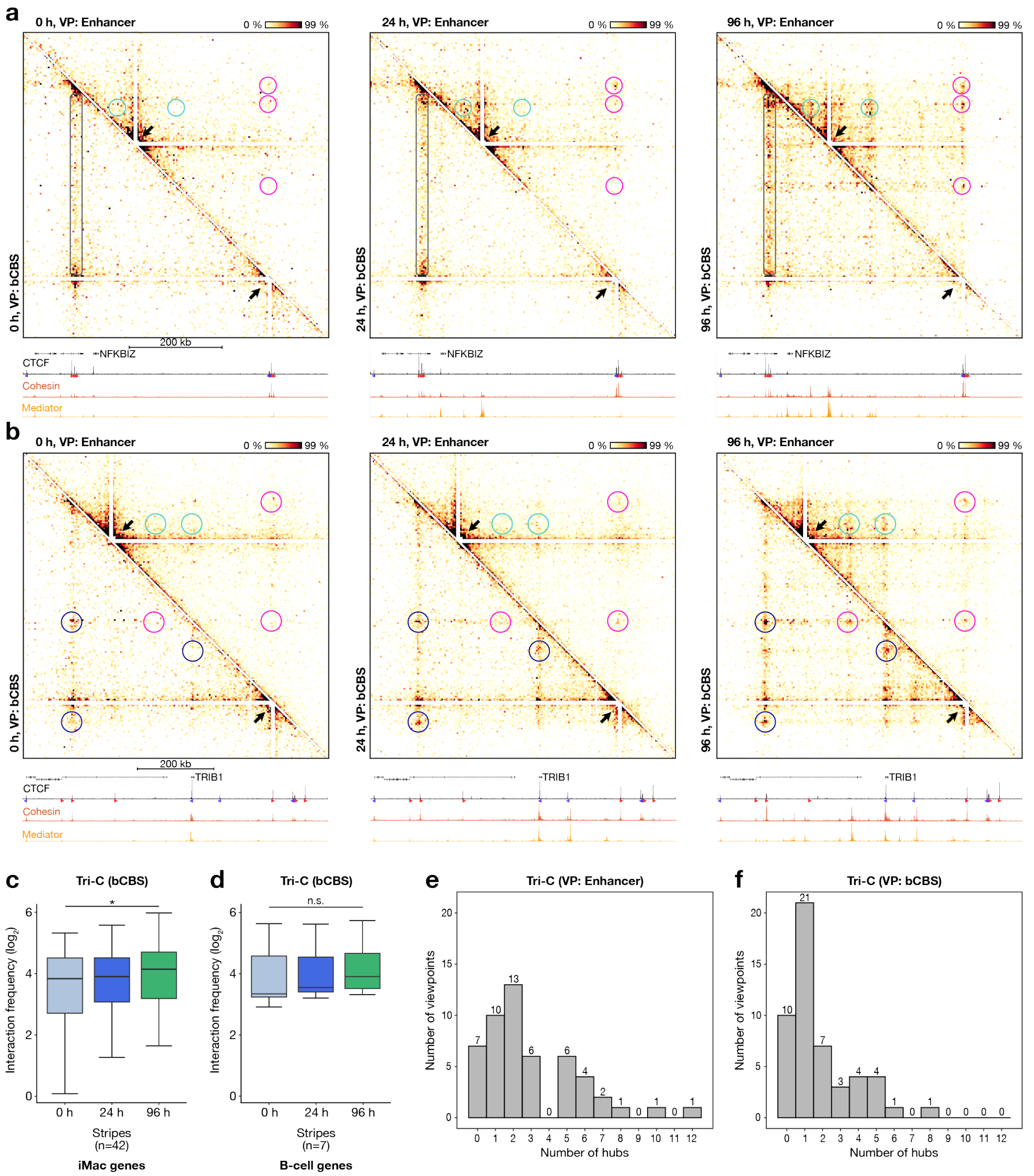


***Extended Data Fig. 3: Dynamic chromatin hub formation during lymphoid-to-myeloid transdifferentiation.*** *(a) Tri-C contact matrices of the NFKBIZ locus (chr3:101,699,405-102,350,405; 651 kb; 3.5 kb resolution) during lymphoid-to-myeloid transdifferentiation at 0 h, 24 h, and 96 h. The top-right matrix shows Tri-C data from the viewpoint of an enhancer; the bottom-left matrix shows the viewpoint of a boundary CTCF-binding site (bCBS). The viewpoints are indicated with white triangles. E-E-P, E-C-X, and C-E/P/C-C-C contacts are highlighted in cyan, magenta, and dark blue circles, respectively. CBS stripes are highlighted in grey rectangles. The profiles below the Tri-C contact matrices show occupancy of CTCF (CTCF ChIP-seq), Mediator (MED26 ChIPmentation), and Cohesin (SMC1A ChIPmentation) at corresponding time points. The axes are scaled to signal and have the following ranges: CTCF = 0–5547; Cohesin = 0–1773; Mediator = 0–2252. (b) Tri-C contact matrices of the TRIB1 locus (chr8:124,988,732-125,788,732; 800 kb; 4 kb resolution) during lymphoid-to-myeloid transdifferentiation, as described in panel a. The axes are scaled to signal and have the following ranges: CTCF = 0–12178; Cohesin = 0–2129; Mediator = 0–5726. (c) Multi-way interaction frequencies of stripes involving a CBS viewpoint and an interacting CBS in iMac-specific loci at 0 h, 24 h, and 96 h after differentiation induction. Boxplots show the interquartile range (IQR) and median of the data; whiskers indicate the minima and maxima within 1.5 * IQR; asterisks indicate significance (P < 0.01, two-sided paired Wilcoxon signed rank test). (d) Multi-way interaction frequencies of CBS stripes in B-cell-specific loci, as described in panel c. (e) Histogram showing the distribution of the number of multi-way interactions with the enhancer viewpoints per locus as detected by Tri-C. (f) Histogram*

*showing the distribution of the number of multi-way interactions with the bCBS viewpoints per locus as detected by Tri-C.*

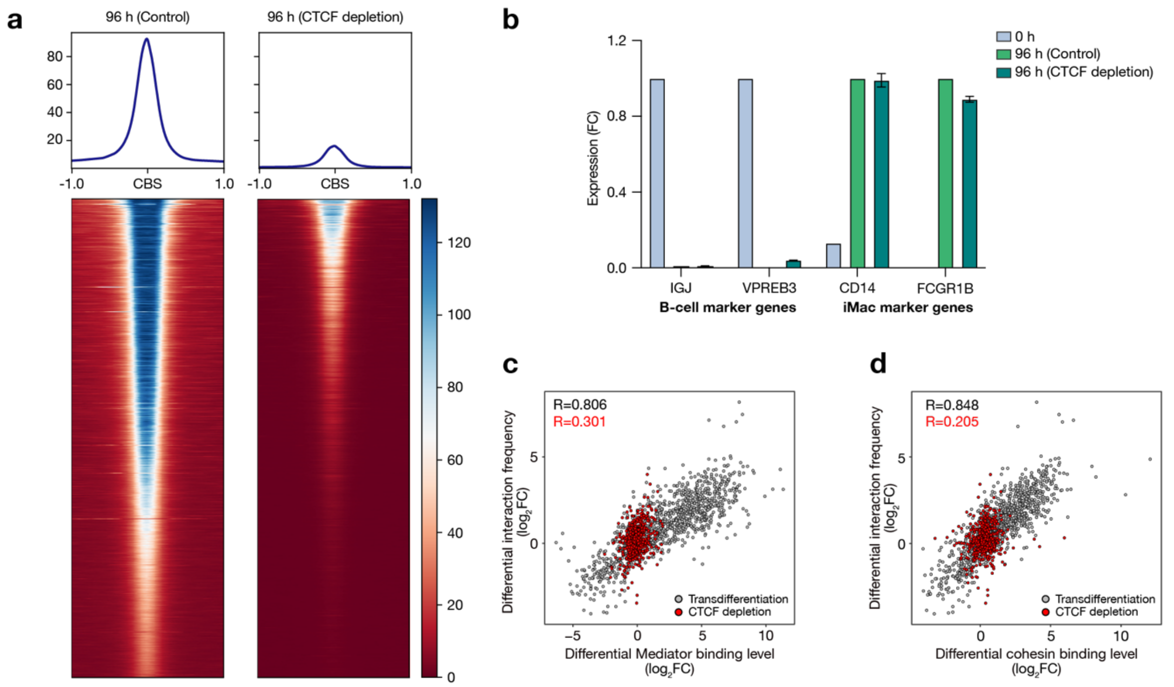


***Extended Data Fig. 4: Characterization of CTCF depletion during lymphoid-to-myeloid transdifferentiation.*** *(a) Comparison of CTCF occupancy in control-treated and CTCF-depleted cells at 96 h after differentiation induction, measured by ChIPmentation. The data show heatmaps of ± 1 kb regions surrounding CTCF-binding sites (CBSs). Merged data of 2 biological replicates are shown. (b) Changes in the expression levels of B-cell and iMac marker genes during lymphoid-to-myeloid transdifferentiation, measured with RT-qPCR at 0 h and at 96 h in control-treated and CTCF-depleted cells. The expression of B-cell marker genes (IGJ, VPREB3) is shown as fold change (FC) relative to the 0 h timepoint, whereas the expression of iMac marker genes (CD14, FCGR1B) is shown relative to 168 h. The bars represent the average of n = 2 replicates, except for the 0 h condition, in which n = 1; the error bars indicate the standard deviation. (c) Correlation between differential enhancer-promoter interaction frequencies and differential Mediator binding levels at the interacting elements during lymphoid-to-myeloid transdifferentiation (grey datapoints; 24 h vs 0h and 96 h vs 0 h) and upon CTCF depletion (red datapoints; 96 h control vs 96 h CTCF depletion), based on Spearman's correlation test. (d) Correlation between differential enhancer-promoter interaction frequencies and differential cohesin binding levels at the interacting elements, as described in panel c.*

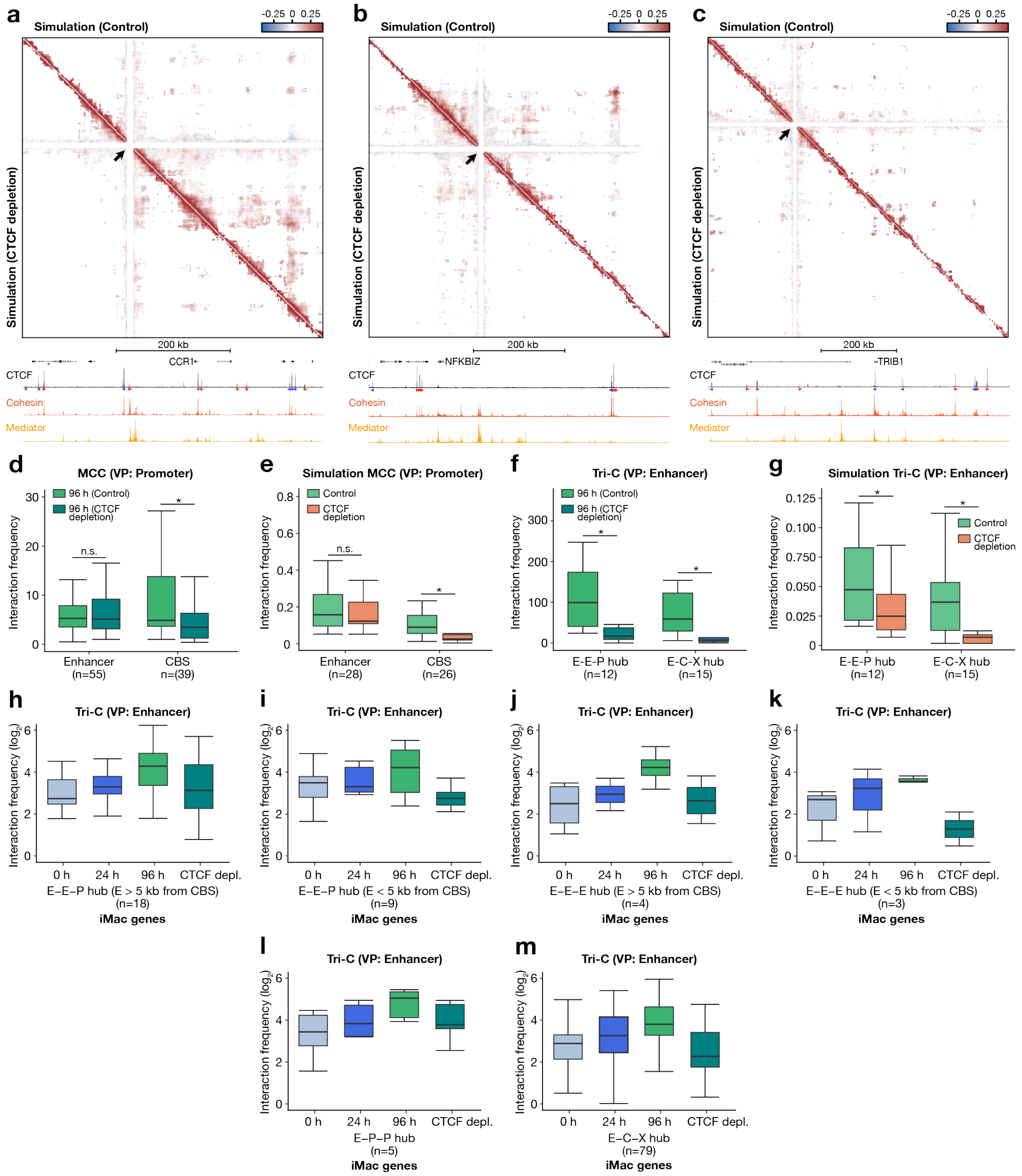


***Extended Data Fig. 5: CTCF supports the formation of enriched multi-way interactions in chromatin hubs.*** *(a) Triplet correlation coefficients of multi-way interactions with the enhancer viewpoint, generated with molecular dynamics simulations of the CCR1 locus (chr3:45,902,299-46,427,299; 525 kb; 2 kb resolution) in control (top-right matrix) and CTCF-depleted (bottom-left matrix) conditions at 96 h after differentiation induction. Correlated (red) regions show a higher propensity for cooperative interactions with the viewpoint than expected based on their pair-wise interaction frequencies (white). The profiles below show occupancy of CTCF (CTCF ChIP-seq), Mediator (MED26 ChIPmentation), and cohesin (SMC1A ChIPmentation) at 96 h. The axes of the profiles are scaled to signal and have the following ranges: CTCF = 0–10069; Cohesin = 0–2960; Mediator = 0–6075. (b) Triplet correlation coefficients of multi-way interactions with the enhancer viewpoint in the NFKBIZ locus (chr3:101,699,405-102,350,405; 651 kb; 2 kb resolution) in control and CTCF-depleted cells at 96 h, as described in panel a. The axes of the profiles are scaled to signal and have the following ranges: CTCF = 0–5547; Cohesin = 0–1773; Mediator = 0–2252. (c) Triplet correlation coefficients of multi-way interactions with the enhancer viewpoint in the TRIB1 locus (chr8:124,988,732-125,788,732; 800 kb; 2 kb resolution) in control and CTCF-depleted cells at 96 h, as described in panel a. The axes are scaled to signal and have the following ranges: CTCF = 0–12178; Cohesin = 0–2129; Mediator = 0–5726. (d) Interaction frequencies of the promoters of modelled gene loci (CCR1, NFKBIZ, and TRIB1) with enhancers and CBSs at 96 h in control and CTCF-depleted cells, derived*

*from experimental MCC data. Boxplots show the interquartile range (IQR) and median of the data; whiskers indicate the minima and maxima within 1.5 * IQR; asterisks indicate significance ($P < 0.01$, two-sided paired Wilcoxon signed rank test). (e) Interaction frequencies of the promoters of modelled gene loci (CCR1, NFKBIZ, and TRIB1) with enhancers and CBSs at 96 h in control and CTCF-depleted cells, extracted from the models. Boxplots as described in panel d. (f) Multi-way interaction frequencies of E-E-P and E-C-X hubs in the modelled gene loci at 96 h in control and CTCF-depleted cells, derived from experimental Tri-C data. Boxplots as described in panel d. (g) Multi-way interaction frequencies of E-E-P and E-C-X hubs in the modelled gene loci at 96 h in control and CTCF-depleted cells, extracted from the models. Boxplots as described in panel d. (h-m) Frequencies of three-way interactions involving two enhancers and a promoter (E-E-P hubs), three-way interactions involving three enhancers (E-E-E hubs), three-way interactions involving one enhancer and two promoters (E-P-P hubs), and three-way interactions involving an enhancer, CTCF-binding site, and any other cis-regulatory element (E-C-X hubs) in iMac-specific loci at 0 h and 24 h and at 96 h in control and CTCF-depleted cells. Boxplots as described in panel d.*

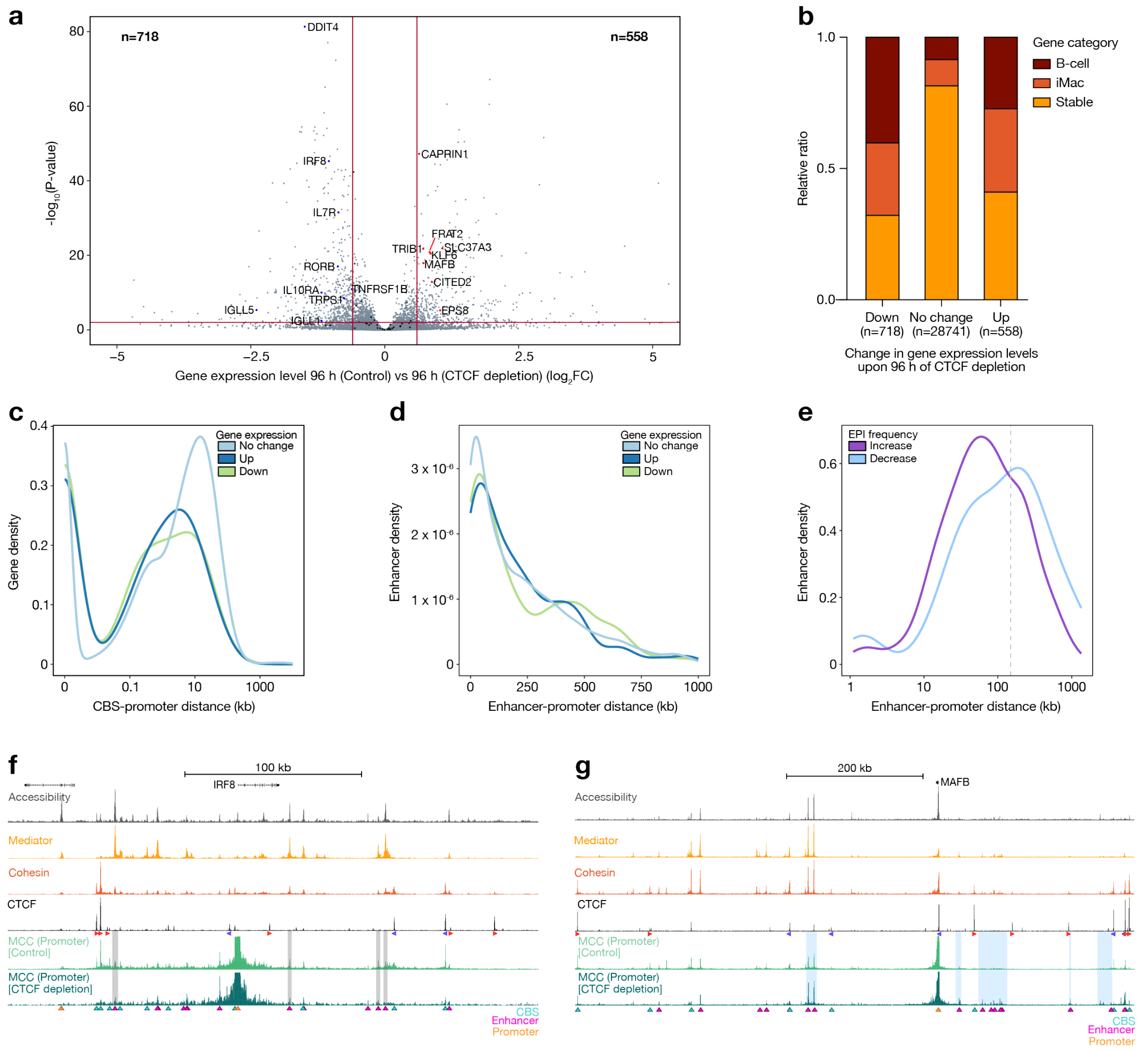


***Extended Data Fig. 6: Changes in gene expression following CTCF depletion can be explained by rewired pair-wise enhancer-promoter interactions.*** *(a) Volcano plot showing differentially expressed genes between control-treated and CTCF-depleted cells at 96 h after differentiation induction, as measured by RNA-seq in n = 2 replicates. The x-axis shows the fold change (FC) in expression; the y-axis shows the adjusted P-value. The horizontal line and vertical lines indicate the significance threshold of adjusted P-value < 0.01 and effect size threshold of $log_2FC > 0.6$ or < -0.6, respectively. Targeted genes are highlighted and those that are significantly changed upon CTCF depletion are labelled. Targeted genes described in the text that are not labelled (e.g., CCR1 and NFKBIZ) are not significantly up- or down-regulated upon CTCF depletion. (b) Comparison of the proportion of B-cell-specific genes, iMac-specific genes, and genes that are stably expressed during lymphoid-to-myeloid-transdifferentiation among the genes that are downregulated, unchanged, or upregulated upon CTCF depletion. (c) Distribution of CBS-promoter distances (of the nearest CBS) of genes that are unchanged, upregulated, or downregulated upon CTCF depletion. (d) Distribution of enhancer-promoter distances (of all paired enhancers, see Methods) of genes that are unchanged, upregulated, or downregulated upon CTCF depletion. (e) Distribution of enhancer-promoter distances of increased and decreased enhancer-promoter interactions upon CTCF depletion in targeted iMac-specific loci as identified by MCC. The grey line marks the 150 kb threshold used to classify distal enhancers in Extended Data Fig. 2c. (f) Chromatin interactions in the IRF8 locus (chr16:85,769,160-86,069,160; 300 kb) in control and CTCF-depleted cells at 96 h after differentiation induction. From top to bottom: gene annotation; chromatin accessibility (ATAC-seq); Mediator occupancy (MED26 ChIPmentation); Cohesin occupancy (SMC1A ChIPmentation); CTCF occupancy (CTCF ChIP-seq); Micro-Capture-C (MCC) data from the viewpoint of the promoter. The axes of the profiles are scaled to signal and have the following ranges: Accessibility = 0–1776; Mediator = 0–4261; Cohesin = 0–3315; CTCF = 0–12417; MCC = 0–40. The orientations of CTCF motifs at prominent CBSs are indicated by arrowheads (forward orientation in red; reverse orientation in blue). MCC interactions with CBSs, enhancers, and promoters are annotated with cyan, magenta, and orange triangles, respectively, and prominent changes in CTCF-depleted cells are highlighted in grey. (g) Chromatin interactions in the MAFB locus (chr20:40,156,606-40,976,606; 820 kb) in control and CTCF-depleted cells, as described in panel*

*f, with prominent changes highlighted in blue. The axes of the profiles are scaled to signal and have the following ranges: Accessibility = 0–3567; Mediator = 0–5231; Cohesin = 0–2232; CTCF = 0–10212; MCC = 0–30.*
